# Human-in-the-loop approach to identify functionally important residues of proteins from literature

**DOI:** 10.1101/2024.03.09.583700

**Authors:** Melanie Vollmar, Santosh Tirunagari, Deborah Harrus, David Armstrong, Romana Gaborova, Deepti Gupta, Marcelo Querino Lima Afonso, Genevieve Evans, Sameer Velankar

## Abstract

We present a novel system that leverages curators in the loop to develop a dataset and model for detecting residue-level functional annotations and other protein structure features from standard publication text. Our approach involves the integration of data from multiple resources, including PDBe, EuropePMC, PubMedCentral, and PubMed, combined with annotation guidelines from UniProt, while employing LitSuggest and Huggingface models as tools in the annotation process. A team of seven annotators manually curated ten articles for named entities, which we utilized to train a starting PubmedBert model from Huggingface. Using a human-in-the-loop annotation system, we developed the best model with commendable performance metrics of 0.90 for precision, 0.92 for recall, and 0.91 for F1-measure.

Our proposed system showcases a successful synergy of machine learning techniques and human expertise in curating a dataset for residue-level functional annotations and protein structure features. The results demonstrate the potential for broader applications in protein research, bridging the gap between advanced machine learning models and the indispensable insights of domain experts.

## Background & Summary

The three-dimensional (3D) structure of a protein determines its function and provides insights into its mechanisms and processes within a cell. In order to understand biology and its intricate systems, it is essential to determine protein structures, analyze them on a residue level and identify which residues are the key to its function. For more than 50 years the Protein Data Bank (PDB) managed by the wwPDB partners^1^ (https://www.wwpdb.org/) has been the go-to data resource to access experimentally determined protein structures. The team at Protein Data Bank in Europe (PDBe)^2^ (https://www.ebi.ac.uk/pdbe/), as one of the founding partners in the wwPDB, processes and curates a couple of hundred new, experimentally derived structure submissions every week. In a unified process, followed by all the wwPDB data centers, they provide a standard set of annotations^3^ alongside the atomic coordinates for each structure. However, the structures are deposited before the publication is available, which prevents biocurators from accessing additional knowledge hidden in scientific literature to support and enrich the protein structure data.

To better understand the structure-function relationship for a protein and its relevance in biological context, it would be beneficial to access additional knowledge locked away in unstructured text in scientific publications. For over 20 years, UniProt^4^ (https://www.uniprot.org/) has developed processes to manually curate scientific literature and enrich protein sequences, the linear, one-dimensional representation of a protein. However, with a near-exponential increase in publication rate and sequences released, it is impossible to comprehensively extract residue-level functional knowledge from literature at scale and annotate structures or sequences solely through manual curation.

Here, we present a workflow to develop a transformer-based named entity recognition (NER) system to annotate full-text publications as the first step to an automated pipeline to extract residue-level functional information from the literature to provide 3D-structure based protein annotations. The presented algorithm achieved high overall precision, recall and F1-measure scores of 0.90, 0.92 and 0.91, respectively.

Identified annotations can be used to highlight key text spans in publications but can also serve as a starting point for future developments. Entity types and their relationships can be collated across all publications for a particular protein. Applying reasoning and weighing to the identified information may help to discover new knowledge and insight into the intricate systems a protein is involved in within cells and organisms, such as new interaction partners or signaling and metabolic pathways, and how these systems are dependent on a protein’s structure-function relationship. Furthermore, these annotations can also serve as a validation source for assessing the biological relevance of predicted protein models. These predicted models are generated by deep learning algorithms such as AlphaFold^5^ and RosettaFold^6^. AlphaFold Protein Structure Database (AlphaFold DB)^7^ (https://alphafold.ebi.ac.uk/) contains predicted models provided by DeepMind (https://deepmind.google/) while other computationally created models can be deposited to the ModelArchive^8^ (https://modelarchive.org/). Large-scale structure predictions can also be made accessible by establishing a 3D-Beacon and integrating it into the 3D-Beacons network^9^ (https://www.ebi.ac.uk/pdbe/pdbe-kb/3dbeacons/) which is designed to improve FAIRness^10^ (Findability, Accessibility, Interoperability and Reusability) of experimental and predicted structure models. Nowadays, the computationally generated models achieve similar quality for chemical and physical descriptors, such as geometry and bond lengths and angles, to those found for experimentally determined structures, at least in areas with high-confidence predictions. Unlike structures in the PDB, these predictions are not supported by experimental evidence like electron density maps or electric potential maps as determined through X-ray crystallography or cryo-electron microscopy, respectively, or chemical shifts from nuclear magnetic resonance. Consequently, analyzing a predicted protein model in isolation is prone to misinterpretation of a residue’s location and its interactions with neighbors. However, non-structural publications can provide information on a variety of biochemical and mutational studies which can be used to check if the predicted models are supported by the observations drawn from these studies. If predicted models can explain the observations, the predicted models could be considered functionally validated.

## Methods

### The annotation team

#### Project manager

The project manager was the lead for the annotation project with more than 15 years of experience in structural biology, more than 500 protein structures in the PDB, and over seven years experience in developing software and machine learning algorithms. As lead, the project manager was responsible for the general management, planning and documentation of the project and was involved in the annotation process.

#### Annotators

A team of six PDBe biocurators, involved in the curation of protein structures submitted to the PDB, volunteered in the annotation process. All but one had a PhD, either in biochemistry, bioinformatics or structural biology with a strong background in biochemistry and/or structural biology. Combined, they had 10 years of experience in bioinformatics, 24 years in biochemistry and structural biology, and 31 years in biocuration. While undertaking the annotation process, the team was split over two different sites and time zones, and annotation was carried out in a fully remote setting.

### Literature selection

The general workflow for literature selection is depicted in Figure 1. In the first step we retrieved all the PubMed (https:// pubmed.ncbi.nlm.nih.gov/) IDs (PMIDs) for publications linked to a protein structure by querying PDBe’s ORACLE database on 29th September 2022. On that date, the PDB contained 196,012 PDB entries with 73,019 associated, unique PMIDs.

**Figure 1.**
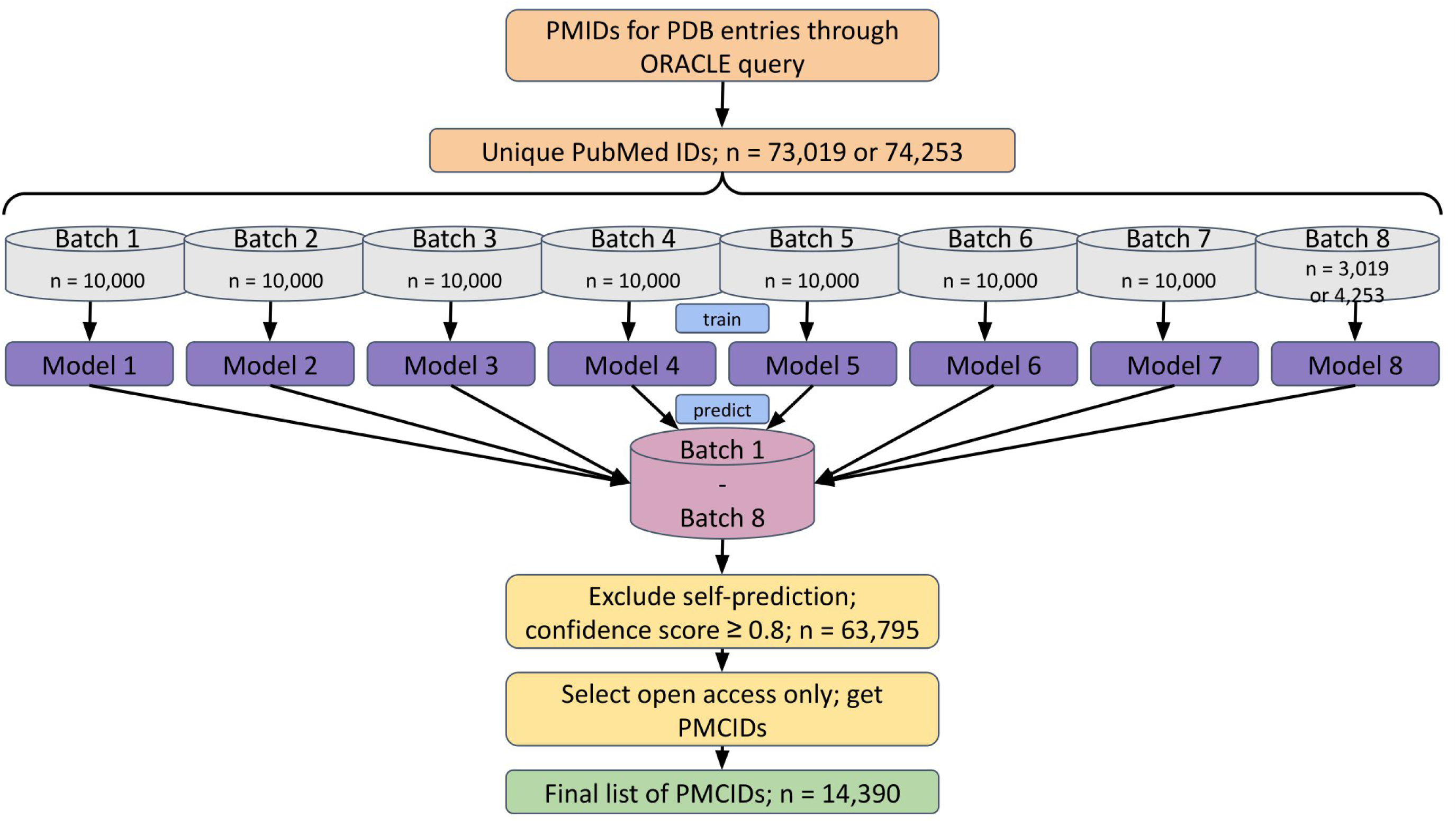
Schema of literature selection workflow.

LitSuggest^11^ (https://www.ncbi.nlm.nih.gov/research/litsuggest/), an AI-driven web browser-based trainable system that directly uses PMIDs, was used as a content filtering tool for assessing the abstract and title of our short-listed publications. Using the list of PMIDs generated above, we created seven publication batches of 10,000 IDs each (the positive samples) and batch 8, an exception, had only 3,019 IDs. All batches were matched with an equal number of randomly picked PMIDs from the entire set in PubMed representing the negative samples. It has to be noted, that due to the selection process, there is a small chance that the negative sets may contain some of the PMIDs from the positive batches. For each batch, a model was trained using the corresponding titles and abstracts for the individual PMIDs in the batch. The trained models were used to identify the relevance of newly added IDs to PubMed over several weeks.

The same exercise was repeated on the 23rd of January 2023, with 200,612 PDB entries having 74,253 unique PMIDs. The additional PMIDs were added to batch 8 which now contained 4,253 IDs while preserving the original batches used in previous training. Each of the eight trained models were presented with 7 batches, excluding the one that was used to train the model to obtain relevance scores for individual PMIDs. A publication was deemed relevant when the predicted confidence score was *≥* 0.8 across all seven cross-prediction models. This resulted in 63,795 (86%) PMIDs predicted as relevant. The prediction statistics across the eight different models are given in Table 1.

**Table 1.** Proportion of publications selected from batches of 10,000 using seven independent LitSuggest models and a confidence score *≥* 0.8.

| batch | number of positive PubMed IDs | percent of total |
| --- | --- | --- |
| 1-10,000 | 8,682 | 87% |
| 10,001-20,000 | 8,626 | 87% |
| 20,001-30,000 | 8,612 | 86% |
| 30,001-40,000 | 8,543 | 85% |
| 40,001-50,000 | 8,601 | 86% |
| 50,001-60,000 | 8,552 | 86% |
| 60,001-70,000 | 8,543 | 85% |
| 70,001-74,253 | 3,580 | 84% |

To adhere to open data principles and to be able to annotate full-text articles, only open access publications with Pub-MedCentral^12^ (https://www.ncbi.nlm.nih.gov/pmc/) IDs (PMCIDs) were identified using the EuropePMC’s^13^ (https://europepmc.org/) article API and included in further works. This further reduced the number of publications included in the study to 14,390, 19% of the initial starting set of 74,253.

Lastly, a number of documents were further rejected during the annotation stage due to a primary focus other than a protein structure, often covering drug and fragment screening campaigns or nucleic acid structures.

### The annotation tool and schema

For our text annotation project, a number of free and paid-for annotation tools were evaluated regarding the following features: compatibility with PubMed and PubMedCentral, possibility for project management, multi-user co-annotation option, integration of ontologies, open source distribution, web browser based, ease of use, and available documentation. The annotation tool of choice for our project was TeamTat^14^ (https://www.teamtat.org/). TeamTat is a free tool and its developers focused on biomedical literature and compatibility with PubMed and PubMedCentral, retrieving publication abstract and metadata such as title, authors, journal and publication year for a given PMID. For open access publications with a PMCID, TeamTat retrieved the entire publication from a BioC XML FTP-server^15^ (https://www.ncbi.nlm.nih.gov/ pmc/tools/openftlist/). The BioC XML format^16^ (https://bioc.sourceforge.net/) was introduced by the BioCreative Initiative^17^ as a way of making scientific publications interoperable. In the case of open access full-text documents, these came with in-line figures, figure captions, tables and table captions.

TeamTat also allowed for project management with the project manager being able to upload/retrieve the relevant literature, assign publications to annotators and control the start and end of an annotation round. Entity types, relationship types and ontology referencing were set up and updated by the project manager. TeamTat also supported versioning and after each annotation round, merging statistics were calculated across the corpus as well as inter-annotator agreement and a new version for the publication set was created. Documents could be exported at any point in the annotation process as either BioC XML or BioC JSON. We opted for the BioC XML format, as it enclosed the plain paragraph text and its identified annotations under the same XML tag (<PASSAGE>), which allowed for easy retrieval of individual sentences with their respective in-line annotations for downstream transformer training.

TeamTat provides access to Medical Subject Headings (MeSH)^18^ and the Gene Ontology (GO)^19,20^ through hard-coded links. Additional ontologies relevant for our project were Sequence Ontology (SO)^21^, Chemical Entities of Biological Interest (ChEBI)^22^, Gene^23^ and PRotein Ontology (PR/PRO)^24^. For each ontology a short-hand name similar to, e.g. “MESH:” for MeSH, was created and served as a prefix to link an entity type to an ontology. A “DUMMY:” short-hand name was used to collect terms that were not found in any of the other ontologies. Although we linked the different entity types to ontologies, controlled vocabularies and reference databases, we did not apply grounding of terms in the annotation process by linking text spans to unique references.

The annotation handbook published by the TeamTat developers (Supplemental Materials of Islamaj et *al*.^25^) was adapted to suit our project requirements. The final detailed annotation schema can be found in Supplemental Material. The project manager generated an initial set of annotated publications to define a set of entity types which formed the basis for developing initial guidelines, which were revisited in the subsequent annotation rounds following discussions with the biocurators (see below in Manual annotation of initial set of publications). The updates to guidelines included adding or removing entity types or clarification on the guidelines. The guidelines continued to be adapted even after switching from fully manual annotation with a team of biocurators to a semi-automated process using a trained model to accommodate the increasingly diverse set of publications. All alterations were done after consultation with the volunteer team of biocurators either in form of open discussion or polling. Focusing on structure and sequence features curated by the UniProt biocurators, we selected entity types that captured details about a particular protein, its structural make-up down to residue level, interaction partners, bound molecules, general properties of the protein, changes to its sequence, organism of origin, experimental methods and evidence to support drawn conclusions. The final list of entity types later used in transformer training was: “bond_interaction”, “chemical”, “complex_assembly”, “evidence”, “experimental_method”, “gene”, “mutant”, “oligomeric_state”, “protein”, “protein_state”, “protein_type”, “ptm”, “residue_name”, “residue_name_number”, “residue_number”, “residue_range”, “site”, “species”, “structure_element”, “taxonomy_domain”. The “Materials and Methods” and “References” sections were excluded from the annotation process as little to no contextual, residue-level information was expected to be present in these sections.

We also developed a detailed user guide (see Supplemental Materials) on how to set up and operate TeamTat from a project manager as well as biocurator perspective. This was used to support the biocurators after initial training when annotating independently.

### Manual annotation of initial set of publications

Initially, ten publications (PMC4784909, PMC4786784, PMC4792962, PMC4832331, PMC4833862, PMC4848090, PMC4850273, PMC4850288, PMC4852598, PMC4887326) were chosen randomly from the filtered, open access list described in the Litera-ture selection section above. Each biocurator was given two publications to manually annotate, based on the guidance from the example annotations and the handbook. A set of two-hour hackathons were organized weekly to annotate the assigned publications. In case the biocurators were not able to attend the hackathon, web-based access through a personalized web-link for the assigned documents was provided to annotate documents outside of the dedicated sessions. The project manager annotated those publications that could not be annotated by the biocurators in order to achieve double-annotation for each document. We acknowledge and are fully aware that the overrepresentation of the project manager annotated documents increased the likelihood of bias. However, even with the best annotation guidelines shaped by a team of expert annotators, assigning entity types to terms is a highly subjective process. A different team of experienced annotators may introduce a different set of biases, based on their training and understanding. The first round of independent annotation lasted approximately four months, after which the annotations across all ten publications were combined and annotation statistics were calculated within TeamTat.

To increase efficiency and accelerate the annotation process, the decision was made to switch from a fully manual to a semi-automated annotation process. The project manager was made responsible for cleaning and consolidating the annotations for the ten initial publications. Upon completion of this task, the cleaned publications were passed to the lead biocurator, who served as a proof-reader. In this capacity, the lead biocurator flagged annotations and entity types that were still ambiguous. In a number of discussions between the project manager and the lead biocurator those ambiguities were resolved and entity types and annotation guidelines were updated. A graphical illustration of the manual annotation workflow can be found in Figure 2. The project manager then applied a final pass of cleaning and consolidating across the ten initial publications before using the annotated text to train a named entity recognition system. This final, consolidated version was used as ground truth against which the annotation performance of each annotator could be measured.

**Figure 2.**
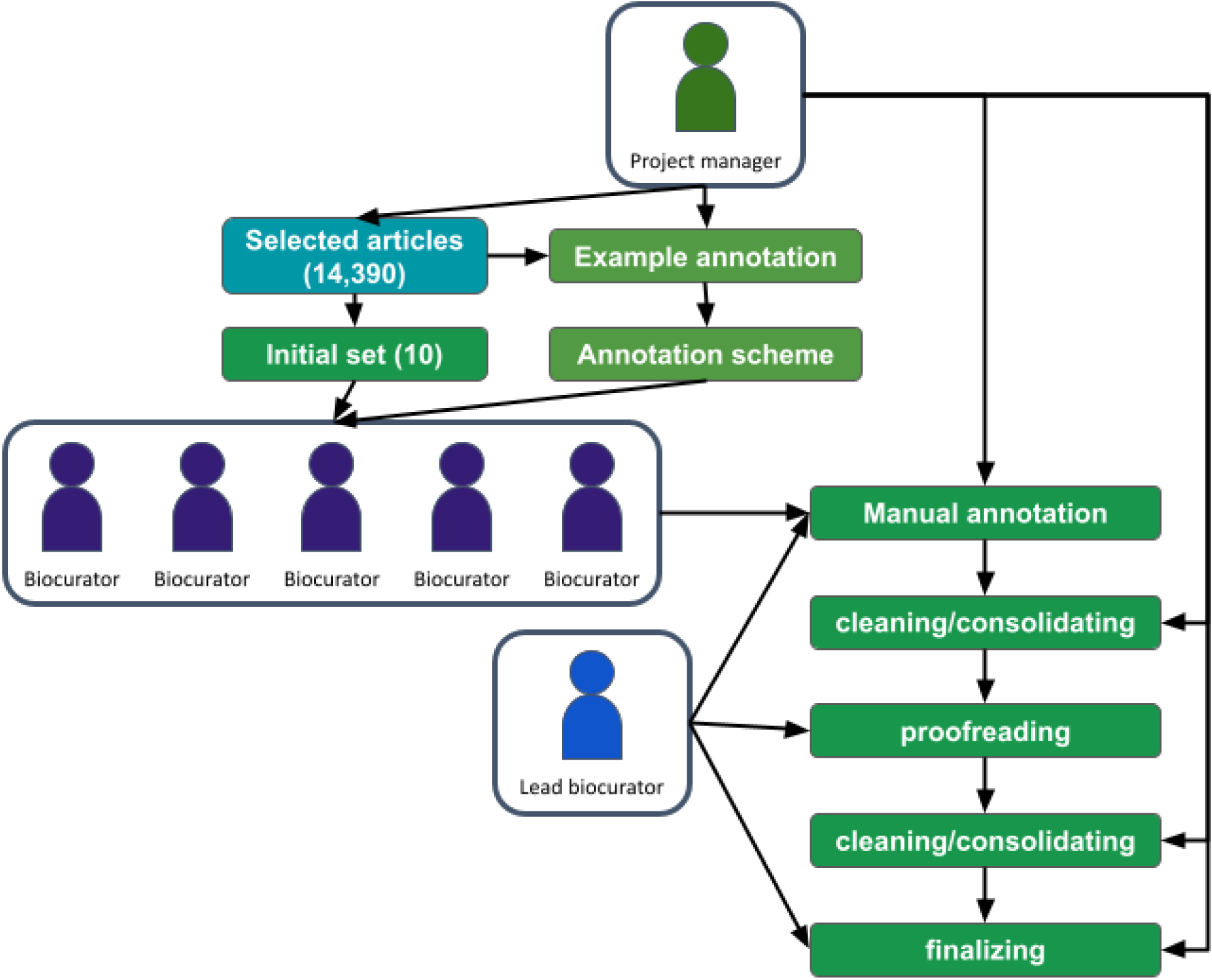
Schema of manual annotation workflow.

### Annotation evaluation

The quality of manual annotations created by the biocurators was judged using the built-in calculation procedures in TeamTat, which follow a partial agreement model. The following six categories of annotation outcomes were determined by TeamTat:

- *FA* - Full Agree: same type, concept ID and text span
- *CA* - Concept Agree: same concept ID and text span, but different types
- *TA* - Type Agree: same type and text span, but different concept IDs
- *PA* - Partial Agree: same type and concept ID for overlapping text
- *DA* - Disagree: different types, different concept IDs for text spans
- *SN* - Single: text annotated by only one of the annotators

The full set of outcomes was only relevant for the initial manual annotation by the biocurators and during the cleaning and remediation steps to create the training data for the initial model. As mentioned in Annotation tool and schema, we did not use concept IDs for grounding terms and only evaluated for prefix matches, which, as they were directly linked to an entity type, always returned a perfect match.

In order to investigate whether there was any bias introduced into the annotations by individual biocurators we also applied the SemEval procedure to the manually annotated publications, see Annotation evaluation using SemEval procedure.

### Annotation processing for training and evaluation

In order to train a transformer-based annotation algorithm and to be able to calculate annotation statistics to monitor the performance of both the algorithm and the human annotators, the publication text and its in-line annotations needed to be converted from BioC XML into the IOB (Inside Outside Beginning) format^26^. For each document we iterated over the individual paragraphs, split them into sentences and combined them with their respective annotations using the offset values available in the BioC XML file. For the total list of isolated sentences, we then generated an index. Next, the isolated sentences were converted into tab-separated TSV files. These TSV files were used to calculate various statistics, see in section Annotation evaluation using SemEval procedure. The index was then randomly split to create three smaller files holding train, test and development sets, containing 70%, 15%, and 15% of sentences, respectively.

During the conversion process it was found that a number of open access documents retrieved from NCBI’s FTP site had line breaks introduced within a paragraph, often in figure or table captions. These line breaks resulted in shifts of the paragraph offset by “+1”, which introduced character position miss-matches for the corresponding annotations. Through personal communication with the maintainers of the FTP site, it was found that the offset shift was likely a result of the conversion process from a number of input file formats provided by publishers to BioC XML. Occasionally, we also found identical sentences and annotations more than once. In this case, only the first occurrence would be included in the data. Although all efforts were made to catch as many errors as possible, on average 21 annotations were lost in each batch, which amounts to 0.2% as each batch had on average 10,037 annotations.

### Training a first named entity recognition system

Using the TSV files from above, we trained a first model. The basic principles of our algorithm and training process are given in Algorithm 1. The training routine described also provided the basis for the iterative training to build a semi-automatic annotation system.

#### Algorithm 1: Iterative Deep Learning Model Training with Curators in Loop

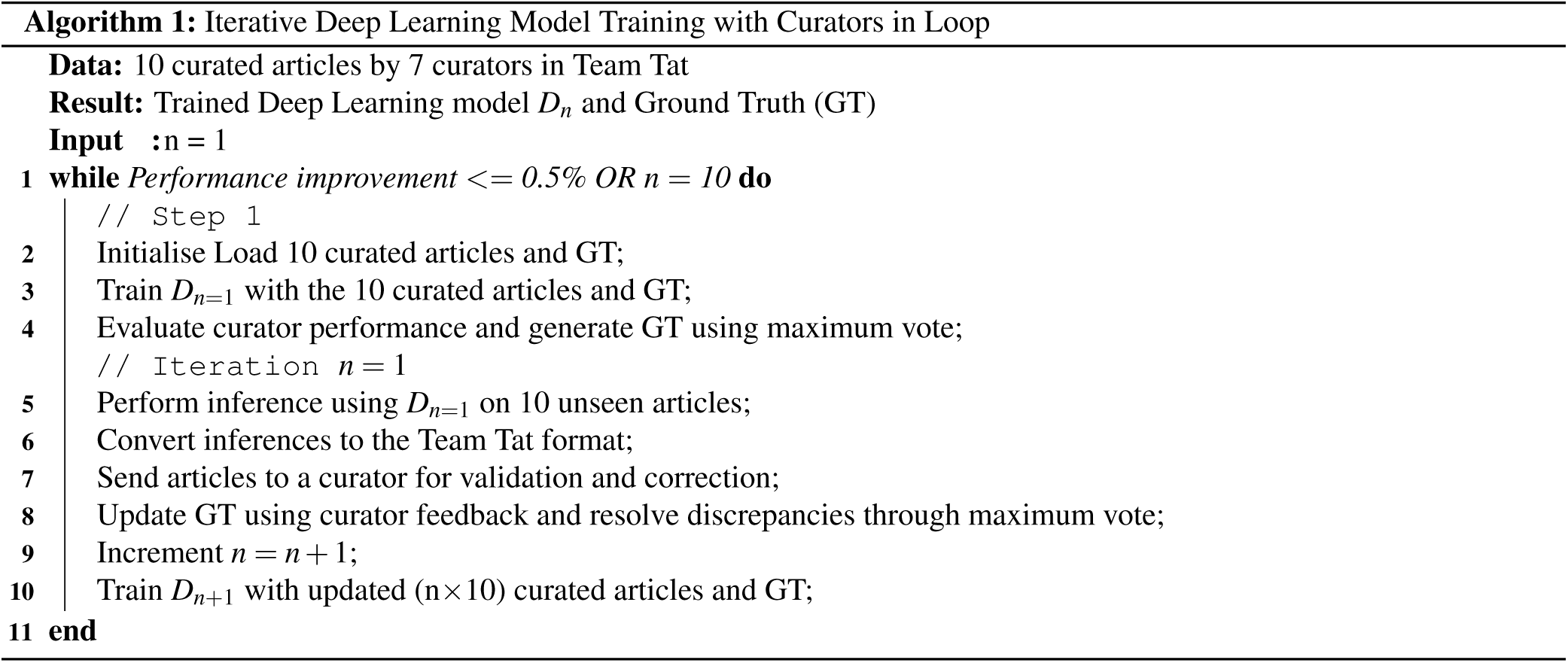

Taking advantage of the rapid developments in natural language processing (NLP), we chose a starting model based on Google’s transformer^27^. For our objective, NER, we looked at BERT (Bidirectional Encoder Representations from Transformers)-based models such as BioBERT^28^, PubmedBERT^29^, and BioFormer^30^. We employed a pre-trained transformer model from Hugging Face (https://huggingface.co), namely *microsoft/BiomedNLP-PubMedBERT-base-uncased-abstract-fulltext*. Fine-tuning was conducted for 3 epochs, initially, using the carefully selected hyperparameters listed in Table 2. Optimizing the hyperparameters resulted in an improved initial model, v1.2, which was used to annotate a new batch of publications. We also reduced the number of entity types from 23 to 19, as we found during the data preparation step that some entity types had too few samples to allow for a meaningful split into train, test and evaluation set and had a negligible contribution to training.

**Table 2.** Hyperparameters for the Named Entity Recognition Model - all models.

| Hyperparameter | Value | Description |
| --- | --- | --- |
| Evaluation Strategy | Epoch-based evaluation | The approach used to evaluate the model during training. |
| Learning Rate | 2e-5 | The step size at which the model's weights are updated during training. |
| Per-Device Training Batch Size | 2 (v1.2, v1.4); 5 (v2.1); 10 (v3.1) | The number of samples used for each gradient update during training on each device. |
| Per-Device Evaluation Batch Size | 2 (v1.2, v1.4); 5 (v2.1); 10 (v3.1) | The number of samples used for evaluation on each device. |
| Number of Training Epochs | 10 | The number of times the entire training dataset is passed through the model during training. |
| Weight Decay | 0.01 | A regularization term used to prevent overfitting by penalizing large weights in the model. |

### Consecutive rounds of semi-automatic annotation and NER training

To develop a robust algorithm, a diverse corpus, larger than the initial ten publications, was needed. Therefore, a human-in-the loop approach combined with a named entity recognition system (see Training a first named entity recognition system) was used, to iteratively increase the number of annotated publications in the corpus.

In each iteration, a new batch of ten publications randomly selected from the open access list was presented to the current best model to identify text spans and annotate them with their entity types. The returned predictions were in BioC XML format which allowed for visual inspection in the annotation tool TeamTat.

At the end of each prediction round, the project manager inspected each of the ten publications in the batch, and fixed any errors in the annotations produced by the NER model. This curation process did not only look at the predicted annotations but rather the pre-annotated spans served as a guide for the annotator, who was still required to read the full text and add missing annotations. Such an approach of post-prediction curation has been implemented as a standard tool in the NLP suite “prodigy” version 3 https://prodi.gy/ in the function “ner.correct” (“ner.make-gold” in version 2). A similar approach was also used by Gnehm et *al*.^31^. Any annotations predicted in the “Methods” and “References” sections were removed (see annotation schema in the Supplemental Materials for details). The curated annotations were stored in BioC XML to be later combined with other batches and converted to the IOB format for a new round of model training or being used as ground truth for model performance monitoring. The applied workflow is presented in Figure 3.

**Figure 3.**
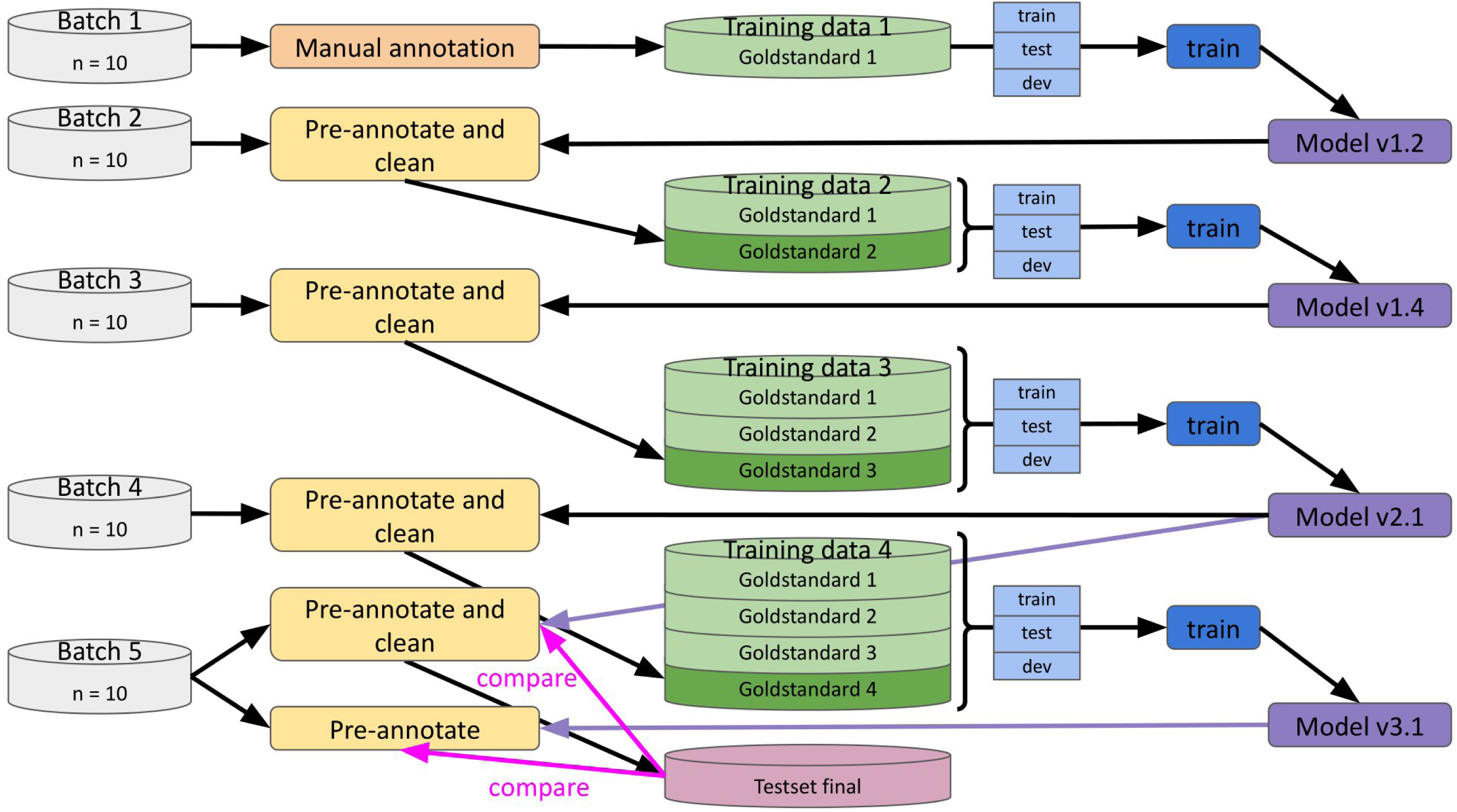
Iterative, human-in-the-loop build up of training data.

For entity types that repeatedly produced large numbers of false-positives or false-negatives, i.e. were not correctly identified by the predictor and required manual curation, anonymous biocurator polling was used to improve the annotation process. Here, examples of ambiguously labeled terms were given to all biocurators and they were asked to assign entity types. A majority vote across the responses determined the entity type and the annotations in the publications were updated accordingly. For example, it was not clear what should be labeled as entity type “*mutant*”. After polling, point mutations at specific sequence positions and deletions/insertions of sequence ranges or whole domains and proteins were included.

The decision was made that no new publications would be added for training once the NER model would not improve by more than 0.5% for overall values for F1-measure, precision and recall across all entity types or reaching 100 publications, whichever came first.

As a result of changes to the annotation schema in terms of entity types (described in section The annotation tool and schema) and adding new publications over time, splitting into train, test and validation sets was carried out anew for every new model training. In order to be able to judge and compare the performance of models v2.1 and v3.1, we therefore employed an additional set of ten publications which had not been used for any training, testing or validation and provided a completely independent test set.

It should also be noted that for inference on a new document, we supplied the publication text split into paragraphs rather than sentences as was used during training. This was aimed at the transformer model’s ability to contextualize named entities and as was shown by Luoma and Pyysalo^32^ and Wang et *al*.^33^ this was expected to improve the model’s performance.

### Annotation evaluation using SemEval procedure

• To monitor and evaluate the performance of the trained predictor, we followed the published assessment process for SemEval^34^. Each predicted annotation was assessed whether it had a matching annotation in the ground truth using the following five categories:

- *Correct* - full agreement between predicted annotation and ground truth annotation in text span and entity type
- *Incorrect* - disagreement between predicted annotation and ground truth annotation in text span and entity type
- *Partial* - text span overlaps in predicted annotation and ground truth annotation but the entity type may differ
- *Missing* - annotation is only found in the ground truth but not in the predicted annotations
- *Spurious* - annotation is only found in the predicted annotations but not in the ground truth SemEval then evaluated a found match whether it belonged to one of four different classes of matches:
- *Strict* - only evaluate annotations with exact text span boundaries and exact entity types between predicted and ground truth annotations
- *Exact* - allow annotations to have exact text span boundaries with disagreement in entity type between predicted and ground truth annotations
- *Partial* - allow annotations to have partially overlapping text span boundaries with disagreement in entity type between predicted and ground truth annotations
- *Type* - allow annotations that have some form of agreement between predicted and ground truth annotations

For each class of match the precision, recall and F1-measure were determined. The statistics were calculated for annotations in the selected sections title, abstract, introduction, results, discussion, tables, as well as table and figure captions. In order to apply the SemEval procedure to the annotations, the text and in-line annotations had to be converted from BioC XML to the IOB format as described above in Annotation processing for training and evaluation.

Please note that the evaluation was done by comparing the predictions to the ground truth. However, we used all the predictions produced by the different models on the full-text BioC XML rather than predicting only on the sentences included in the ground truth. As a consequence, the models produced predictions that are not found in the ground truth. Those additional sentences were given the O(utside) label and appended to the ground truth. During the evaluation process these annotations were classed as *“spurious”*.

For the batches used during consecutive training described above in Consecutive rounds of semi-automatic annotation and NER training, we performed the evaluation across the entire batch. The independent test set, batch 5, for the comparison of autoannotator versions v2.1 and v3.1, additionally underwent a per-document evaluation.

## Results

### Manual annotation statistics and annotator performance

After merging the annotations from the initial manual annotation round, it became apparent that only seven out of ten publications had been double-annotated, namely PMC4784909, PMC4786784, PMC4792962, PMC4832331, PMC4833862, PMC4850273, and PMC4852598. The remaining three publications had very few double or only single annotations. We therefore calculated inter-annotator agreement using the TeamTat analysis tool for both options, seven documents with double annotations and ten documents with three only having partial annotations. The statistics are provided in Table 3. Excluding the partially annotated publications caused a drop of 1.8% for the full-agreement statistics from 79% for ten documents to 77.2% for seven. Although gold-standard double-annotation could not be achieved for all documents, those that were fully annotated had high agreement between the annotators and were expected to provide good quality training data for bootstrapping a transformer model.

**Table 3.**
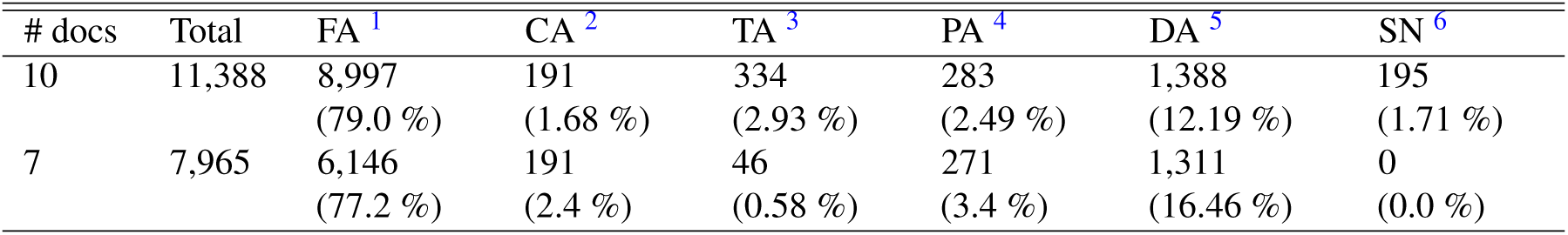
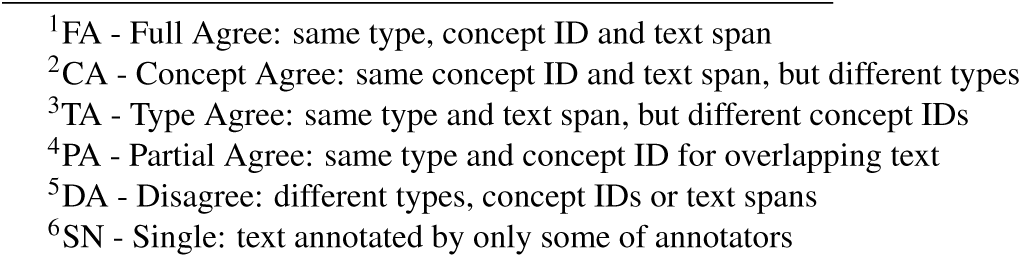
Inter-annotator agreement between the biocurators for the initial ten publications after the first independent annotation round.

| # docs | Total | FA <sup>1</sup> | CA <sup>2</sup> | TA <sup>3</sup> | PA <sup>4</sup> | DA <sup>5</sup> | SN <sup>6</sup> |
| --- | --- | --- | --- | --- | --- | --- | --- |
| 10 | 11,388 | 8,997<br>(79.0 %) | 191<br>(1.68 %) | 334<br>(2.93 %) | 283<br>(2.49 %) | 1,388<br>(12.19 %) | 195<br>(1.71 %) |
| 7 | 7,965 | 6,146<br>(77.2 %) | 191<br>(2.4 %) | 46<br>(0.58 %) | 271<br>(3.4 %) | 1,311<br>(16.46 %) | 0<br>(0.0 %) |

<sup>1</sup>FA - Full Agree: same type, concept ID and text span
<sup>2</sup>CA - Concept Agree: same concept ID and text span, but different types
<sup>3</sup>TA - Type Agree: same type and text span, but different concept IDs
<sup>4</sup>PA - Partial Agree: same type and concept ID for overlapping text
<sup>5</sup>DA - Disagree: different types, concept IDs or text spans
<sup>6</sup>SN - Single: text annotated by only some of annotators

After further cleaning and term disambiguation discussed between the project manager and the lead biocurator, a consolidated set of annotated documents was created, which served as a ground truth. Table 4 gives the total annotation counts for each publication in the ground truth. We manually annotated 10,451 terms, across all publications ranging from 715 to 1,549 and a mean of 1,045 annotations per article. Those annotations represented 2,988 unique terms across 19 different entity types. Table 5 contains the raw counts as well as the number of unique terms found for each entity type. We found that the top-10 entity types for total number of counts were also those with the highest count for unique terms. Those specific entity types sorted by total count in descending order are: *“protein”*, *“structure_element”*, *“protein_state”*, *“chemical”*, *“residue_name_number”*, *“protein_type”*, *“evidence”*, *“mutant”*, *“experimental_method”* and *“site”*. They represent the most relevant key terms to describe residue-level details in a protein structure and do not only appear with high frequency in the training data, but also provide the algorithm with a diverse set of terms to learn and generalize from.

**Table 4.** Total number of annotation count for each of the initial ten manually annotated publications, after curation and consolidation.

| Document ID | annotation count |
| --- | --- |
| PMC4784909 | 865 |
| PMC4786784 | 1,549 |
| PMC4792962 | 1,268 |
| PMC4832331 | 739 |
| PMC4833862 | 1,044 |
| PMC4848090 | 987 |
| PMC4850273 | 1,121 |
| PMC4850288 | 716 |
| PMC4852598 | 1,229 |
| PMC4887326 | 933 |
| Sum | 10,451 |

**Table 5.** Total and unique annotation count for the different entity types in the initial ten manually annotated publications, after curation and consolidation.

| Entity type | Total count | Unique mentions |
| --- | --- | --- |
| Chemical | 1,030 | 260 |
| Complex_assembly | 289 | 81 |
| Evidence | 645 | 155 |
| Experimental_method | 570 | 302 |
| Gene | 154 | 43 |
| Mutant | 614 | 188 |
| Oligomeric_state | 151 | 23 |
| Protein | 1,457 | 122 |
| Protein_state | 1,093 | 303 |
| Protein_type | 756 | 267 |
| PTM | 134 | 40 |
| Residue_name | 139 | 49 |
| Residue_name_number | 795 | 281 |
| Residue_number | 48 | 20 |
| Residue_range | 90 | 85 |
| Site | 519 | 209 |
| Species | 205 | 60 |
| Structure_element | 1,448 | 453 |
| Taxonomy_domain | 314 | 47 |
| Sum | 10,451 | 2,988 |

This ground truth was also used to assess each biocurator’s performance in the initial annotation round using the SemEval evaluation process. The evaluation was done for all the publications for an annotator, regardless of whether they had been fully or partially annotated. In Table 6, the precision achieved by each annotator is given. Applying a “partial agreement” evaluation strategy, we found that all annotators reach a score of 0.79 and above, which underlines the fact that all biocurators have a strong biochemistry and structural biology background and generally look for the same terms within a publication and find most of the occurrences in the ground truth. The scores for the recall, Table 7, are much lower which indicates that all biocurators are not consistent in annotating the same terms, i.e. within the same document a term may have been assigned different entity types, if a term spans multiple words there may be different span boundaries for the same term or a term may have been missed. The F1-measure for each annotator in Table 8 again supports the finding that generally all annotators share the same understanding for the key terms in the documents but differ in their assignment to a specific entity type and where the span boundaries should be placed. It is worth noting that Annotator0 achieved the highest scores for precision, recall and F1-measure for all four evaluation options. Such dominance from one annotator increases the risk of bias. However, considering that the agreement between different annotators, given in Table 3, is *>* 75%, i.e. annotators agree at least partially on more than 75% of annotations, the majority of annotations will not have been biased by Annotator0’s performance.

**Table 6.** Precision for manual annotation compared to ground truth for each annotator using SemEval evaluation.

| annotator | strict | exact | partial | type | # documents |
| --- | --- | --- | --- | --- | --- |
| annotator0 | 0.82 | 0.88 | 0.92 | 0.87 | 8 |
| annotator1 | 0.70 | 0.80 | 0.87 | 0.74 | 2 |
| annotator2 | 0.52 | 0.80 | 0.88 | 0.59 | 2 |
| annotator3 | 0.55 | 0.65 | 0.79 | 0.69 | 2 |
| annotator4 | 0.49 | 0.71 | 0.85 | 0.72 | 2 |
| annotator5 | 0.49 | 0.86 | 0.92 | 0.53 | 2 |
| annotator6 | 0.78 | 0.90 | 0.94 | 0.82 | 2 |

**Table 7.** Recall for manual annotation compared to ground truth for each annotator using SemEval evaluation.

| annotator | strict | exact | partial | type | # documents |
| --- | --- | --- | --- | --- | --- |
| annotator0 | 0.64 | 0.69 | 0.72 | 0.68 | 8 |
| annotator1 | 0.39 | 0.44 | 0.49 | 0.41 | 2 |
| annotator2 | 0.37 | 0.58 | 0.63 | 0.42 | 2 |
| annotator3 | 0.43 | 0.50 | 0.62 | 0.61 | 2 |
| annotator4 | 0.05 | 0.08 | 0.09 | 0.08 | 2 |
| annotator5 | 0.14 | 0.24 | 0.25 | 0.14 | 2 |
| annotator6 | 0.20 | 0.23 | 0.24 | 0.21 | 2 |

**Table 8.** F1-measure for manual annotation compared to ground truth for each annotator using SemEval evaluation.

| annotator | strict | exact | partial | type | # documents |
| --- | --- | --- | --- | --- | --- |
| annotator0 | 0.72 | 0.77 | 0.81 | 0.76 | 8 |
| annotator1 | 0.50 | 0.57 | 0.62 | 0.53 | 2 |
| annotator2 | 0.43 | 0.67 | 0.74 | 0.42 | 2 |
| annotator3 | 0.48 | 0.57 | 0.69 | 0.61 | 2 |
| annotator4 | 0.10 | 0.14 | 0.17 | 0.15 | 2 |
| annotator5 | 0.21 | 0.37 | 0.40 | 0.23 | 2 |
| annotator6 | 0.32 | 0.36 | 0.38 | 0.33 | 2 |

Overall, the annotator statistics underline the high level of expert knowledge of the biocurators and that, although gold standard double annotation was not achieved across all documents and some bias from one annotator may have been introduced, the identified annotations are of high quality.

### Initial model trained on ten manually annotated publications

The initial model (v1.2) was trained on ten manually annotated publications described in Manual annotation of initial set of publications. Its performance results on the development set are plotted in Figure 4. Throughout the training process, the model’s performance consistently improved, as indicated by decreasing losses and increasing precision, recall, F1-measure, and accuracy. However, the increasing gap between training and validation loss indicated overfitting. With the small sample size of 10,451 annotations used to develop this first model, overfitting was not surprising, but learning was clearly observed in the loss curves. Consequently, we increased the sample size and explored some hyperparameter settings and iteratively improved the model (see below Consecutive model training).

**Figure 4.**
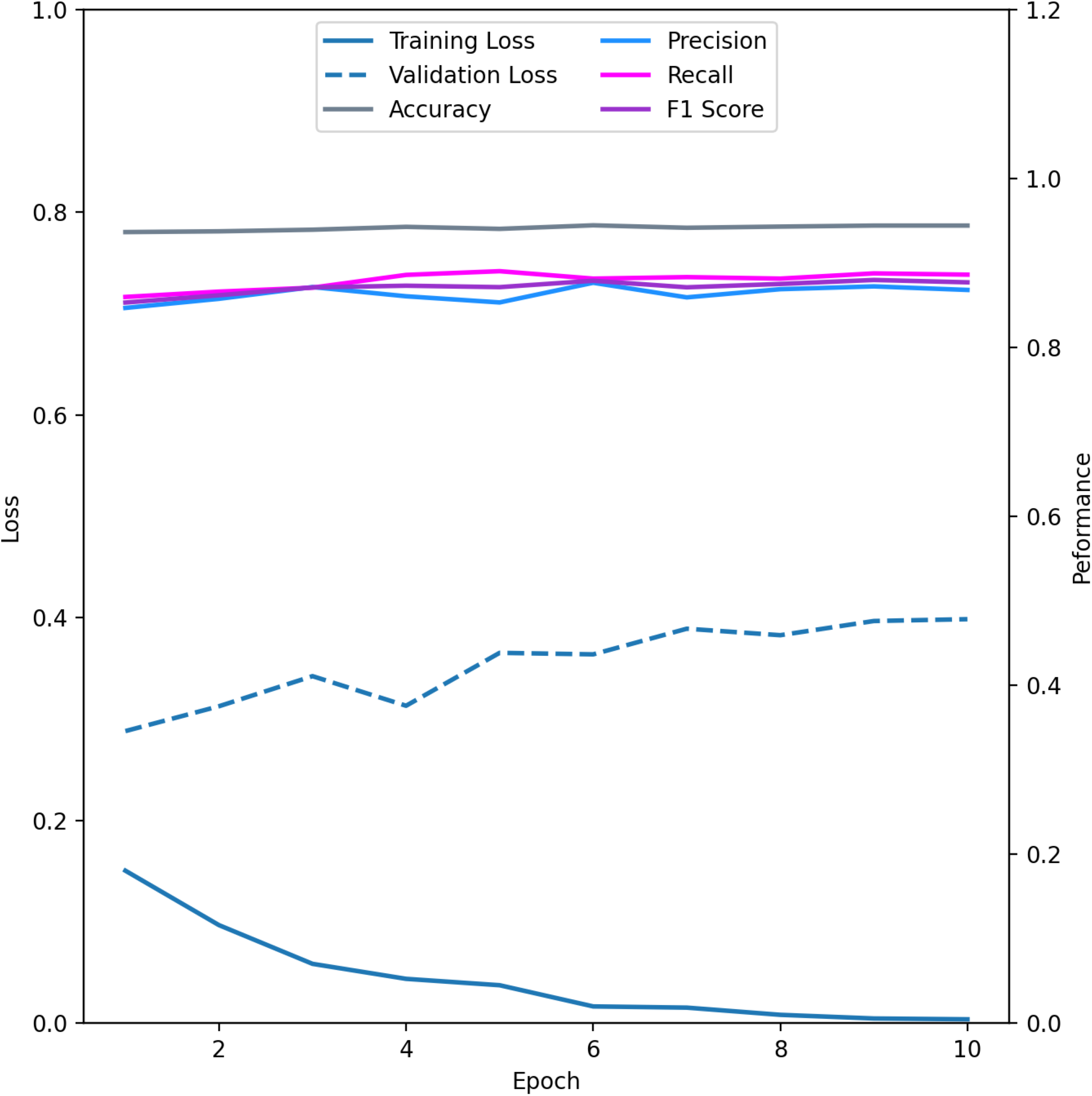
Training and validation loss for the first model v1.2 as well as accuracy, precision, recall and F1-measure.

### Consecutive model training

In Table 9 we give the overall performance statistics (precision, recall, F1-measure and accuracy) for the respective test sets of the different consecutive models averaged across all entity types. Although there is only minimal change in the overall statistics for the different models, plotting the training and validation loss for each training round and comparing the different models revealed that those trained on the larger corpus are less prone to overfitting (see Figure 5). With each additional batch of annotations added, the training loss for the corresponding model started at a higher point, as would be expected from a larger, more diverse corpus, requiring a model to work harder to learn commonalities in the data. This followed a sharp drop, which lasted for the first three epochs and, finally, all models converged to a similar value after seven epochs. Therefore, all models showed clear signs of learning. To assess how the models performed on data not used for training, we looked at the validation loss determined for each respective validation set. For models v1.2 and v1.4 the validation loss never dropped, but instead showed a continuous increase, which is a clear sign of overfitting (see Figure 5). A model that is prone to overfitting is undesirable, as this would lead to unreliable predictions. After adding additional annotations for models v2.1 and v3.1, we observed a similar drop in the validation loss as we found in the training loss (Figure 5) for the first three epochs followed by a slow increase thereafter. Such behavior indicated that the models trained on a larger corpus exhibited reduced overfitting and that predictions for new, unseen data could be expected to be of similar quality as for the training data.

**Figure 5.**
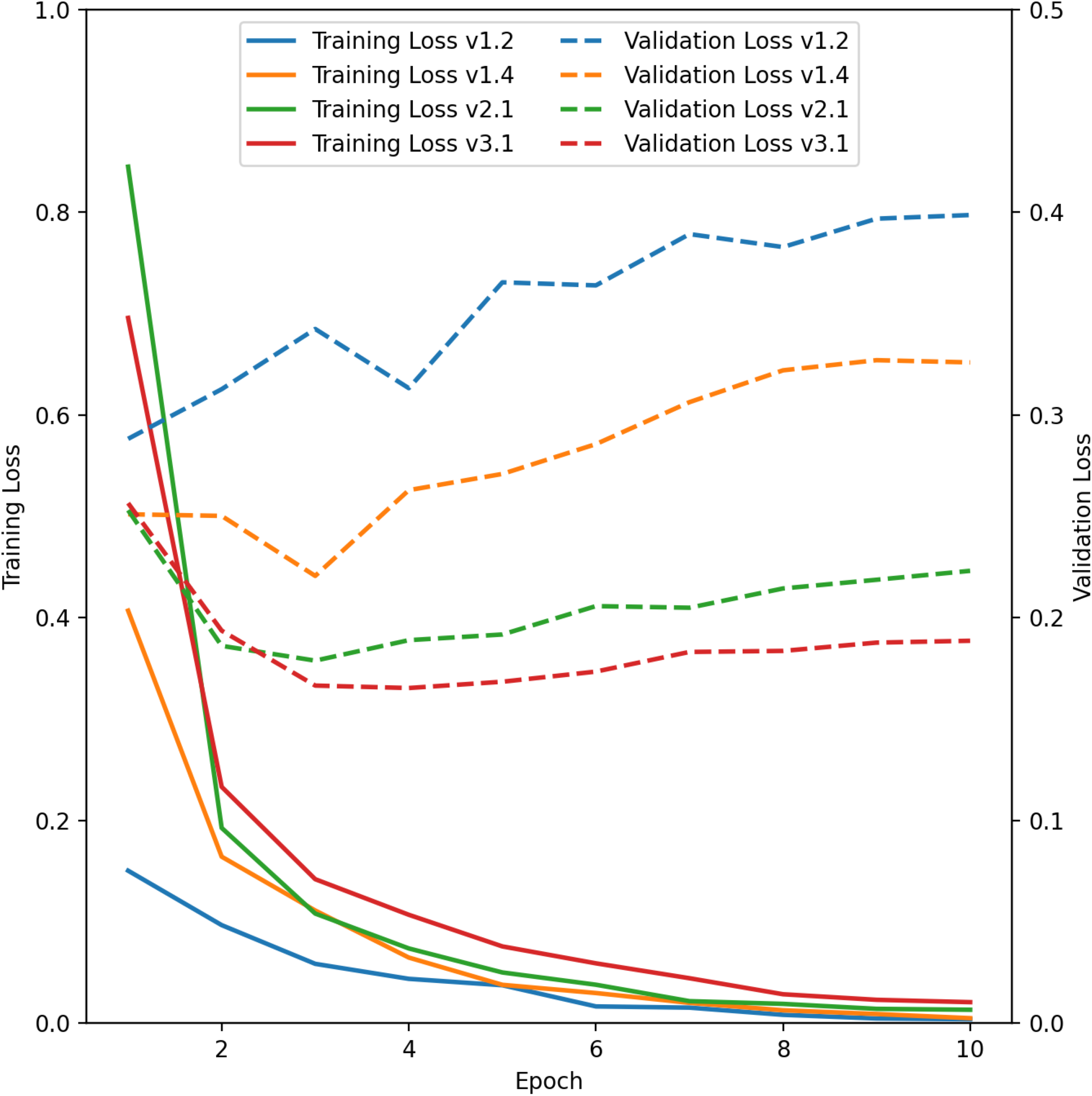
Training and validation loss for the different consecutive models v1.2, v1.4, v2.1 and v3.1.

**Table 9.** Overall training statistics for consecutive models.

| model | Overall precision | Overall recall | Overall F1-measure | Overall accuracy |
| --- | --- | --- | --- | --- |
| v1.2 | 0.87 | 0.89 | 0.88 | 0.95 |
| v1.4 | 0.90 | 0.92 | 0.91 | 0.96 |
| v2.1 | 0.90 | 0.92 | 0.91 | 0.96 |
| v3.1 | 0.91 | 0.92 | 0.91 | 0.96 |

To further judge the performance of the different models, we also monitored the changes in precision, recall and F1-measure for the individual entity types in the respective test sets, Table 10, Table 11, Table 12, respectively. Generally, even the model trained on only ten publications (v1.2) already did reasonably well across the different entity types with the vast majority of scores for precision, recall and F1-measure being *≥* 0.85. However, as observed above in the general training and validation loss plots, model v1.2, and to a lesser degree v1.4, was prone to overfitting, which is supported by achieving scores of 1.00 for a number of entity types for precision, recall and F1-measure. We also found that not all entity types were predicted with similar confidence by the different models. Such behavior reflects the fact that the training data is highly imbalanced for the different entity types, but also that the annotation schema was updated between models. Therefore, a direct comparison of the different models has to be done with caution.

**Table 10.** Precision for the different models for the different entity types on the test set.

| Entity | v1.2 | v1.4 | v2.1 | v3.1 |
| --- | --- | --- | --- | --- |
| Bond Interaction | - | - | 0.93 | 0.82 |
| Chemical | 0.84 | 0.90 | 0.89 | 0.92 |
| Complex Assembly | 0.85 | 0.88 | 0.91 | 0.89 |
| Evidence | 0.74 | 0.86 | 0.84 | 0.89 |
| Experimental Method | 0.77 | 0.73 | 0.85 | 0.80 |
| Gene | 0.86 | 0.89 | 0.79 | 0.79 |
| Mutant | 0.83 | 0.93 | 0.91 | 0.92 |
| Oligomeric State | 0.94 | 0.88 | 0.93 | 0.96 |
| Protein | 0.91 | 0.97 | 0.94 | 0.96 |
| Protein State | 0.80 | 0.78 | 0.83 | 0.86 |
| Protein Type | 0.85 | 0.84 | 0.85 | 0.85 |
| PTM | 0.88 | 0.64 | 0.70 | 0.85 |
| Residue Name | 0.86 | 0.97 | 0.92 | 0.74 |
| Residue Name Number | 0.99 | 0.98 | 0.95 | 0.96 |
| Residue Number | 1.00 | 1.00 | 0.80 | 0.70 |
| Residue Range | 1.00 | 0.86 | 0.81 | 0.89 |
| Site | 0.83 | 0.83 | 0.85 | 0.88 |
| Species | 0.96 | 0.97 | 0.94 | 0.95 |
| Structure Element | 0.88 | 0.91 | 0.91 | 0.91 |
| Taxonomy Domain | 0.95 | 0.97 | 0.99 | 0.98 |

**Table 11.** Recall for the different models for the different entity types on the test set.

| Entity | v1.2 | v1.4 | v2.1 | v3.1 |
| --- | --- | --- | --- | --- |
| Bond Interaction | - | - | 0.88 | 0.91 |
| Chemical | 0.90 | 0.93 | 0.91 | 0.91 |
| Complex Assembly | 0.76 | 0.91 | 0.93 | 0.90 |
| Evidence | 0.76 | 0.89 | 0.88 | 0.88 |
| Experimental Method | 0.75 | 0.76 | 0.85 | 0.82 |
| Gene | 0.92 | 0.86 | 0.86 | 0.65 |
| Mutant | 0.92 | 0.95 | 0.97 | 0.94 |
| Oligomeric State | 1.00 | 1.00 | 0.99 | 1.00 |
| Protein | 0.93 | 0.97 | 0.97 | 0.96 |
| Protein State | 0.83 | 0.85 | 0.88 | 0.88 |
| Protein Type | 0.84 | 0.90 | 0.85 | 0.88 |
| PTM | 0.76 | 0.81 | 0.70 | 0.79 |
| Residue Name | 0.95 | 0.92 | 0.97 | 0.96 |
| Residue Name Number | 0.99 | 0.99 | 0.96 | 0.98 |
| Residue Number | 1.00 | 0.93 | 0.97 | 0.73 |
| Residue Range | 0.80 | 0.91 | 0.70 | 0.86 |
| Site | 0.82 | 0.86 | 0.87 | 0.90 |
| Species | 0.98 | 1.00 | 0.96 | 0.95 |
| Structure Element | 0.86 | 0.92 | 0.92 | 0.92 |
| Taxonomy Domain | 0.97 | 0.96 | 0.98 | 0.98 |

**Table 12.**
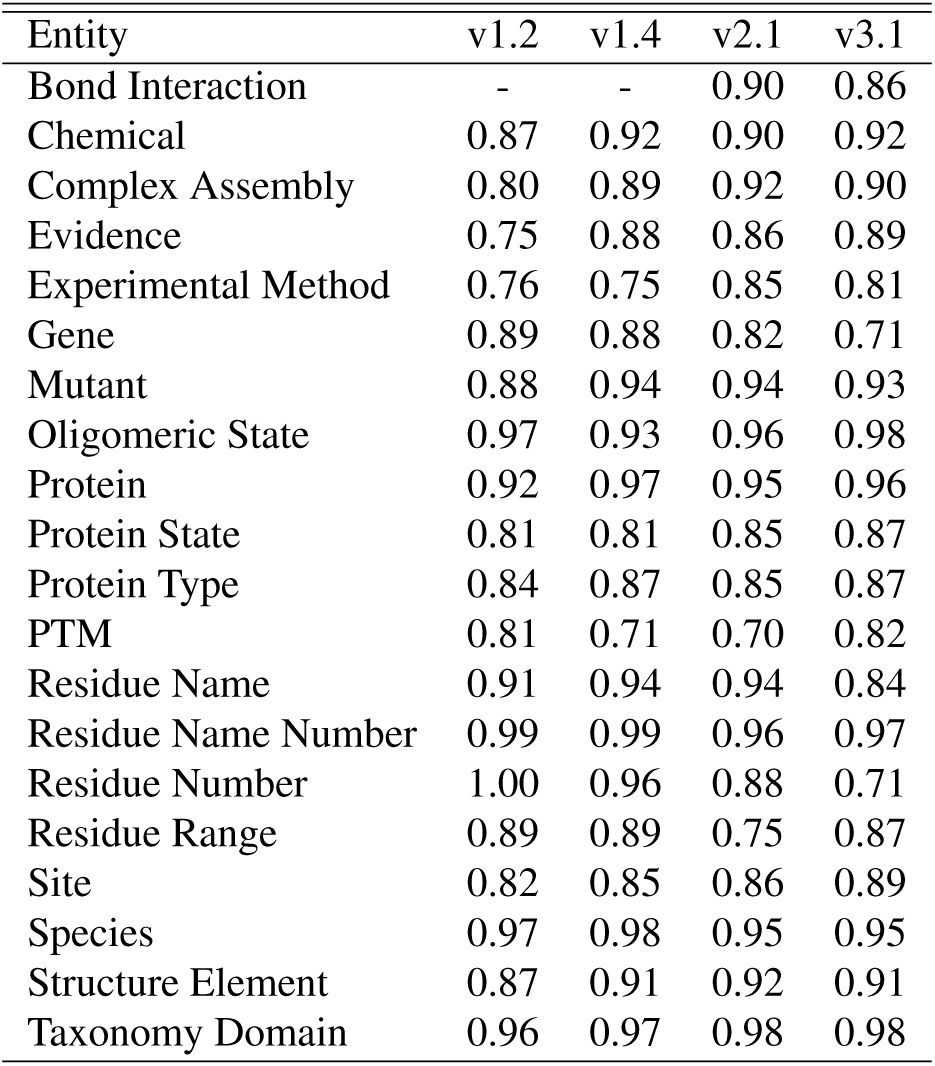
F1-measure for the different models for the different entity types on the test set.

| Entity | v1.2 | v1.4 | v2.1 | v3.1 |
| --- | --- | --- | --- | --- |
| Bond Interaction | - | - | 0.90 | 0.86 |
| Chemical | 0.87 | 0.92 | 0.90 | 0.92 |
| Complex Assembly | 0.80 | 0.89 | 0.92 | 0.90 |
| Evidence | 0.75 | 0.88 | 0.86 | 0.89 |
| Experimental Method | 0.76 | 0.75 | 0.85 | 0.81 |
| Gene | 0.89 | 0.88 | 0.82 | 0.71 |
| Mutant | 0.88 | 0.94 | 0.94 | 0.93 |
| Oligomeric State | 0.97 | 0.93 | 0.96 | 0.98 |
| Protein | 0.92 | 0.97 | 0.95 | 0.96 |
| Protein State | 0.81 | 0.81 | 0.85 | 0.87 |
| Protein Type | 0.84 | 0.87 | 0.85 | 0.87 |
| PTM | 0.81 | 0.71 | 0.70 | 0.82 |
| Residue Name | 0.91 | 0.94 | 0.94 | 0.84 |
| Residue Name Number | 0.99 | 0.99 | 0.96 | 0.97 |
| Residue Number | 1.00 | 0.96 | 0.88 | 0.71 |
| Residue Range | 0.89 | 0.89 | 0.75 | 0.87 |
| Site | 0.82 | 0.85 | 0.86 | 0.89 |
| Species | 0.97 | 0.98 | 0.95 | 0.95 |
| Structure Element | 0.87 | 0.91 | 0.92 | 0.91 |
| Taxonomy Domain | 0.96 | 0.97 | 0.98 | 0.98 |

The iterative training approach meant that for every batch that was automatically annotated by one of the models we also had a set of ground truth annotations from curation, which served as training data for the next generation model, but could also be used for evaluation of the predecessor. Following the SemEval protocol, we evaluated the performance of the different models. The statistics for precision, recall and F1-measure for each model and its respective ground truth batch are given in tables Table 13, Table 14 and Table 15. We found that with each training iteration using an increased and more diverse set of annotations, the scores for precision, recall and F1-measure improved, reflecting a model’s ability to make predictions closer to the respective ground truth.

**Table 13.** Precision for models and their respective publication batches compared to ground truth for each batch using SemEval evaluation.

| model | data batch | strict | exact | partial | type |
| --- | --- | --- | --- | --- | --- |
| v1.2 | batch 2 | 0.66 | 0.75 | 0.83 | 0.75 |
| v1.4 | batch 3 | 0.69 | 0.79 | 0.85 | 0.75 |
| v2.1 | batch 4 | 0.75 | 0.83 | 0.88 | 0.82 |
| v2.1 | batch 5 | 0.78 | 0.85 | 0.91 | 0.85 |
| v3.1 | batch 5 | 0.77 | 0.84 | 0.90 | 0.86 |

**Table 14.** Recall for models and their respective publication batches compared to ground truth for each batch using SemEval evaluation.

| model | data batch | strict | exact | partial | type |
| --- | --- | --- | --- | --- | --- |
| v1.2 | batch 2 | 0.67 | 0.77 | 0.85 | 0.76 |
| v1.4 | batch 3 | 0.71 | 0.80 | 0.87 | 0.77 |
| v2.1 | batch 4 | 0.75 | 0.83 | 0.89 | 0.83 |
| v2.1 | batch 5 | 0.75 | 0.82 | 0.87 | 0.81 |
| v3.1 | batch 5 | 0.75 | 0.79 | 0.86 | 0.81 |

**Table 15.**
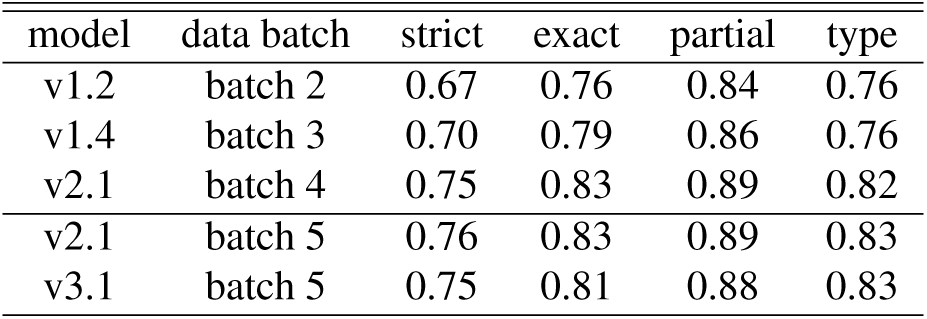
F1-measure for models and their respective publication batches compared to ground truth for each batch using SemEval evaluation.

| model | data batch | strict | exact | partial | type |
| --- | --- | --- | --- | --- | --- |
| v1.2 | batch 2 | 0.67 | 0.76 | 0.84 | 0.76 |
| v1.4 | batch 3 | 0.70 | 0.79 | 0.86 | 0.76 |
| v2.1 | batch 4 | 0.75 | 0.83 | 0.89 | 0.82 |
| v2.1 | batch 5 | 0.76 | 0.83 | 0.89 | 0.83 |
| v3.1 | batch 5 | 0.75 | 0.81 | 0.88 | 0.83 |

Finally, we compared models v2.1 and v3.1 on an independent test set of ten publications to serve as ground truth for final model selection. The prediction statistics for the two models on the individual documents in the independent test set are given in Table 16 for precision, Table 17 for recall and F1-measure in Table 18. The statistics were calculated following the SemEval process. We found that both models performed well on the independent test set with some publications proving easier to predict on, i.e. models achieving higher scores. As model v2.1 achieved higher scores for the different evaluation types across all documents, we chose it as the current best model.

**Table 16.** Precision for models v2.1 and v3.1 on individual documents of batch 5 using SemEval evaluation.

| Document ID | v2.1 |  |  |  | v3.1 |  |  |  |
| --- | --- | --- | --- | --- | --- | --- | --- | --- |
|  | strict | exact | partial | type | strict | exact | partial | type |
| PMC4806292 | 0.78 | 0.84 | 0.91 | 0.87 | 0.78 | 0.84 | 0.91 | 0.86 |
| PMC4817029 | 0.86 | 0.90 | 0.94 | 0.91 | 0.86 | 0.89 | 0.94 | 0.92 |
| PMC4980666 | 0.74 | 0.76 | 0.83 | 0.82 | 0.72 | 0.75 | 0.83 | 0.82 |
| PMC4981400 | 0.74 | 0.81 | 0.89 | 0.87 | 0.76 | 0.83 | 0.89 | 0.86 |
| PMC4993997 | 0.78 | 0.84 | 0.91 | 0.85 | 0.78 | 0.81 | 0.90 | 0.85 |
| PMC5012862 | 0.85 | 0.92 | 0.95 | 0.89 | 0.83 | 0.90 | 0.94 | 0.89 |
| PMC5014086 | 0.74 | 0.87 | 0.93 | 0.83 | 0.72 | 0.84 | 0.92 | 0.83 |
| PMC5063996 | 0.77 | 0.88 | 0.92 | 0.83 | 0.77 | 0.87 | 0.92 | 0.85 |
| PMC5173035 | 0.69 | 0.81 | 0.89 | 0.75 | 0.68 | 0.77 | 0.87 | 0.80 |
| PMC5603727 | 0.74 | 0.80 | 0.89 | 0.85 | 0.74 | 0.79 | 0.88 | 0.86 |

**Table 17.**
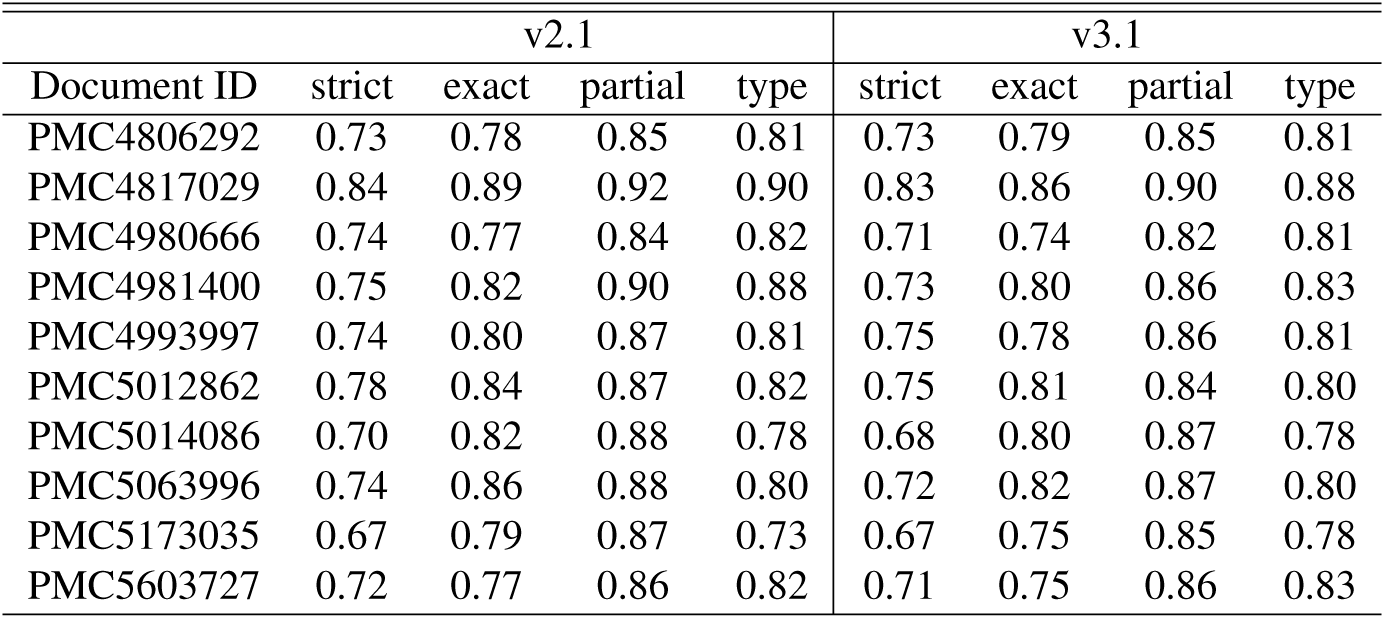
Recall for models v2.1 and v3.1 on individual documents of batch 5 using SemEval evaluation.

| Document ID | v2.1 |  |  |  | v3.1 |  |  |  |
| --- | --- | --- | --- | --- | --- | --- | --- | --- |
|  | strict | exact | partial | type | strict | exact | partial | type |
| PMC4806292 | 0.73 | 0.78 | 0.85 | 0.81 | 0.73 | 0.79 | 0.85 | 0.81 |
| PMC4817029 | 0.84 | 0.89 | 0.92 | 0.90 | 0.83 | 0.86 | 0.90 | 0.88 |
| PMC4980666 | 0.74 | 0.77 | 0.84 | 0.82 | 0.71 | 0.74 | 0.82 | 0.81 |
| PMC4981400 | 0.75 | 0.82 | 0.90 | 0.88 | 0.73 | 0.80 | 0.86 | 0.83 |
| PMC4993997 | 0.74 | 0.80 | 0.87 | 0.81 | 0.75 | 0.78 | 0.86 | 0.81 |
| PMC5012862 | 0.78 | 0.84 | 0.87 | 0.82 | 0.75 | 0.81 | 0.84 | 0.80 |
| PMC5014086 | 0.70 | 0.82 | 0.88 | 0.78 | 0.68 | 0.80 | 0.87 | 0.78 |
| PMC5063996 | 0.74 | 0.86 | 0.88 | 0.80 | 0.72 | 0.82 | 0.87 | 0.80 |
| PMC5173035 | 0.67 | 0.79 | 0.87 | 0.73 | 0.67 | 0.75 | 0.85 | 0.78 |
| PMC5603727 | 0.72 | 0.77 | 0.86 | 0.82 | 0.71 | 0.75 | 0.86 | 0.83 |

**Table 18.**
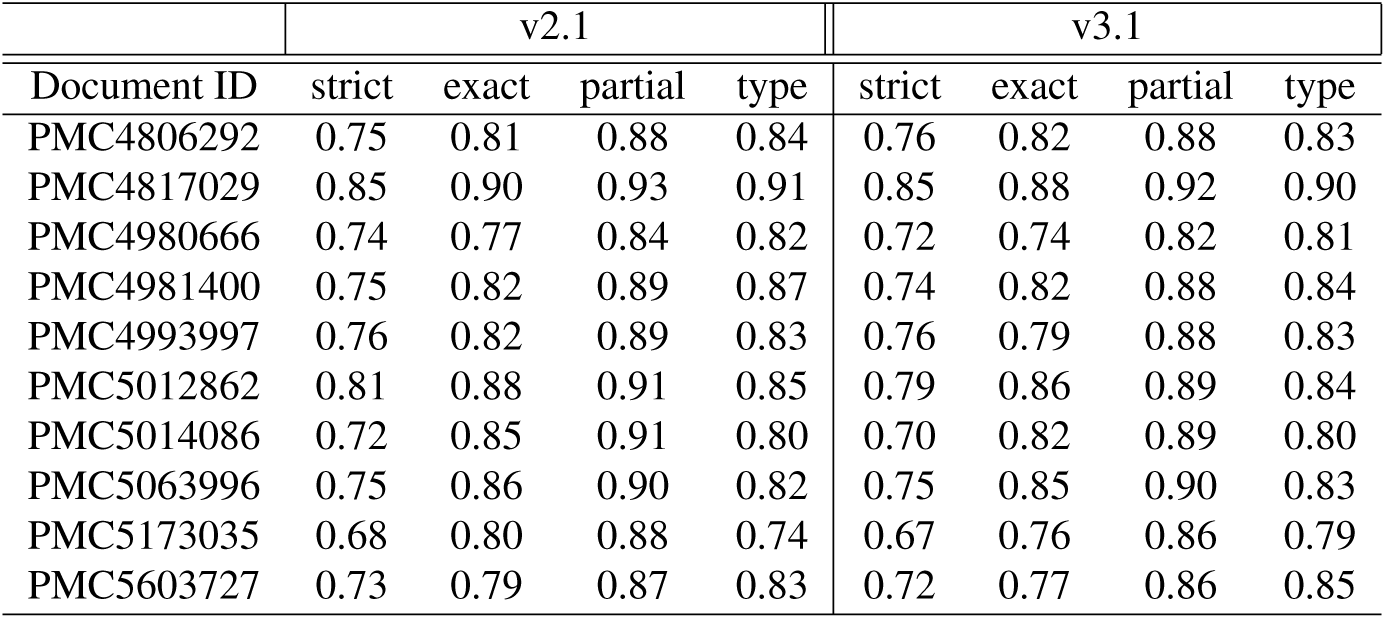
F1-measure for models v2.1 and v3.1 on individual documents of batch 5 using SemEval evaluation.

| Document ID | v2.1 |  |  |  | v3.1 |  |  |  |
| --- | --- | --- | --- | --- | --- | --- | --- | --- |
|  | strict | exact | partial | type | strict | exact | partial | type |
| PMC4806292 | 0.75 | 0.81 | 0.88 | 0.84 | 0.76 | 0.82 | 0.88 | 0.83 |
| PMC4817029 | 0.85 | 0.90 | 0.93 | 0.91 | 0.85 | 0.88 | 0.92 | 0.90 |
| PMC4980666 | 0.74 | 0.77 | 0.84 | 0.82 | 0.72 | 0.74 | 0.82 | 0.81 |
| PMC4981400 | 0.75 | 0.82 | 0.89 | 0.87 | 0.74 | 0.82 | 0.88 | 0.84 |
| PMC4993997 | 0.76 | 0.82 | 0.89 | 0.83 | 0.76 | 0.79 | 0.88 | 0.83 |
| PMC5012862 | 0.81 | 0.88 | 0.91 | 0.85 | 0.79 | 0.86 | 0.89 | 0.84 |
| PMC5014086 | 0.72 | 0.85 | 0.91 | 0.80 | 0.70 | 0.82 | 0.89 | 0.80 |
| PMC5063996 | 0.75 | 0.86 | 0.90 | 0.82 | 0.75 | 0.85 | 0.90 | 0.83 |
| PMC5173035 | 0.68 | 0.80 | 0.88 | 0.74 | 0.67 | 0.76 | 0.86 | 0.79 |
| PMC5603727 | 0.73 | 0.79 | 0.87 | 0.83 | 0.72 | 0.77 | 0.86 | 0.85 |

## Conclusion

Manually annotating a domain specific corpus with a team of experts is usually a time and cost intensive process. Our presented approach shows that a small corpus of ten publications annotated by a team of highly qualified experts is sufficient to bootstrap an initial model, v1.2, for a human-in-the-loop application. In consecutive rounds, we improved on this initial model to yield our best performing version, v2.1. Our best model is highly specific in annotating biomedical literature concerned with protein structures and should be used to identify key-terms describing the 3-dimensional composition of proteins. We made all our data and models open access and they are available for download or programmatic access from https://huggingface.co/. The code provided alongside can be used as a wrapper to run and evaluate the models locally or serve as a start to develop a similar workflow in any other field.

## Data availability

The various data files have been made available at: https://huggingface.co/PDBEurope

1. Stand-alone curator annotations

- CSV
- JSON
- Inside-outside-beginning (IOB)
2. Full-text XML files (without annotations)
3. Full-text BioC with annotations in XML format
4. Full-text BioC with annotations in JSON format

The annotations and documents are made available in a number of formats. We provide the annotations and publications grouped as they were used to train the models v1.2, v1.4, v2.1 and v3.1. The annotations for batch 5, used to compare models v2.1 and v3.1 are provided separately. The plain, full-text XML files of all documents are provided in BioC without annotations. The annotations themselves are provided in-line in BioC for each publication, either as XML or as JavaScript Object Notation (JSON) format. Additionally, the annotations are available as standalone comma-separated values (CSV) and JSON. In these standalone CSV and JSON files, an annotation is described by its unique “*id*”, “*character start*”, “*character end*” to locate the starting and ending character positions within a sentence, “*span*” representing the covered text span and “*entity type*” giving its entity type. To identify from which document an annotation was retrieved, we also use the PMCID of the corresponding publication as “*document id*” in the standalone annotation files. We also provide the annotations and their respective sentences in the IOB format. The IOB files provide sentences with IOB tags and follow the CoNLL NER corpus standards^34^. The datasets to develop the four different models and the independent test set are available from Huggingface:

- https://huggingface.co/datasets/PDBEurope/protein_structure_NER_model_v1.2
- https://huggingface.co/datasets/PDBEurope/protein_structure_NER_model_v1.4
- https://huggingface.co/datasets/PDBEurope/protein_structure_NER_model_v2.1
- https://huggingface.co/datasets/PDBEurope/protein_structure_NER_model_v3.1
- https://huggingface.co/datasets/PDBEurope/protein_structure_NER_independent_val_ set

For example, the data folder for model v1.2 contains the following subfolders and files:

- annotated_BioC_JSON: one JSON file for each annotated publication in BioC; *< PMCID >*_ann.json
- annotated_BioC_XML: one XML file for each annotated publication in BioC; *< PMCID >*_ann.xml
- annotation_CSV: one CSV file for each publication, annotations only; *< PMCID >*.csv
- annotation_IOB: all annotated sentences in IOB format and training, testing, development subsets; all.tsv, train.tsv, test.tsv, dev.tsv
- annotation_JSON: single JSON file containing all annotations from all documents
- raw_BioC_XML: one XML file for each NOT annotated publication in BioC; *< PMCID >*_raw.xml

The four trained models, v1.2, v1.4, v2.1 and v3.1 are available from Huggingface:

- https://huggingface.co/PDBEurope/BiomedNLP-PubMedBERT-ProteinStructure-NER-v1.2
- https://huggingface.co/PDBEurope/BiomedNLP-PubMedBERT-ProteinStructure-NER-v1.4
- https://huggingface.co/PDBEurope/BiomedNLP-PubMedBERT-ProteinStructure-NER-v2.1
- https://huggingface.co/PDBEurope/BiomedNLP-PubMedBERT-ProteinStructure-NER-v3.1

## Code availability

The code is available at the repository https://github.com/PDBeurope/ner_for_protein_structures. Detailed documentation and how to install the tools can be found at https://ner-for-protein-structures. readthedocs.io/en/latest/. Scripts include those used for cleaning and formatting of the annotations from annotation tool TeamTat and how to generate the datasets in the IOB format for input to deep learning algorithms. Additional scripts allow the calculation of model performance and prediction outcome following the SemEval process.

## Acknowledgements

We would like to thank Rezarta Islamaj Doğan (National Center for Biotechnology Information, National Library of Medicine, Bethesda, MD 20894, USA) for her patience to explain aspects of the annotation tool TeamTat to M.V. and give some highly valued tips towards the design of the annotation schema. Funding was provided to M.V. as ARISE Fellowship from the European Union’s Horizon 2020 research and innovation programme under the Marie Skłodowska-Curie grant agreement No 945405.

## Author contributions statement

M.V. identified relevant literature, designed and developed annotation guidelines and handbooks, conceived the experiment(s), developed all the scripts for data preparation and formatting, trained the NER models, analysed the results, generated all the figures and wrote the manuscript. S.T. provided guidance, starting code base and advice for training the NER models and conducting SemEval analysis and contributed to writing the manuscript. D.H. took part in the annotation of the ten initial documents, helped shape the annotation guidelines. D.A., R.G., D.G., M.Q.L.A. and G.E. took part in the annotation of the ten initial documents and reviewed the manuscript. S.V. and D.H. reviewed the manuscript.

