## Supplemental Materials for "Human-in-the-loop approach to identify functionally important residues of proteins from literature"

### Human-in-the-loop NER to identify functionally important residues of proteins from literature - Supplemental material

March 2024

### 1 Annotation handbook

#### 1.1 Entity types

Below are the entity types that were fixed at a corpus size of 30 publications. Some examples are given in Suppl Table 1.

***Bond_interaction***

Any type of covalent or non-covalent bond or interaction. This covers, for example, hydrogen bonds, salt bridges, stacking of aromatic amino acid side chains and nucleic acids as $\pi-\pi$ stacking, electrostatic, hydrophobic, hydrophilic interactions and van der Waals interactions. These interactions are usually created by specific residues at a specific position in the protein of interest. This also covers covalent bonds that have been created between the primary protein of interest and a *"chemical"* as part of a reaction mechanism. Excluded are disulfide bridges as they fall under *"ptm"* and artificially introduced bonds from cross-linking experiments.

***Chemical***

This covers any chemical compound that is found in the publication and is not a protein. Most commonly this refers to small molecules/ drugs/compounds and fragments thereof, co-factors/ligands/metals and other molecules interacting with the primary protein that are not a protein themselves. This also includes short peptides and peptide fragments that may have been used instead of the corresponding full-length protein. Amino and nucleic acids are also included if they represent a substrate or product of a reaction carried out by the primary protein rather than a specific residue in a sequence. Unspecific references such as "moiety" or "group" are excluded.

***Complex_assembly***

A higher order, often heterogeneous, assembly or complex is covered by this entity type. The primary protein of interest serves as the main component in such a complex and interacts with another component, e.g. another *"protein"* or *"chemical"*, and this interaction can be measured, even if only transiently. This is different from the entity type *"oligomeric_state"* as distinct oligomeric states of the primary protein can be combined to higher order complexes.

***Evidence***

Any measured quantity or experimental output that provides support for any conclusion drawn is annotated as *"evidence"*.

***Experimental_method***

This covers any procedure/experiment/method employed to produce the *"evidence"* that supports the conclusions drawn.

***Gene***

This refers to a specific gene/operon/open reading frame mentioned in the publication and is often used as a synonym for the gene product, i.e. the protein of interest.

***Mutant***

This represents a sequence-edited version of the primary protein or a related protein for comparison. The changes covered are point mutations of a specific residue in a specific sequence position and deletion/insertion mutants where whole sections/domains have been removed or added to a protein. We also include domain swaps between proteins to create chimeras. However, alterations such as expression tags, fluorescence labeling or adding other reporter tags are included in *"experimental_method"*.

***Oligomeric_state***

The *"oligomeric_state"* is only concerned with the polypeptide and no other components like those covered by *"chemical.* As mentioned above, this is different from *"complex_assembly"* as some proteins require a distinct oligomeric state, i.e. require multiple copies of the same polypeptide, to be in their active form or even stable. As such, smaller oligomers serve as building blocks that can be assembled into higher order complexes.

***Protein***

The primary protein of interest or related proteins (homologues/orthologues/paralogues from other *"species"*) used for additional analysis and comparison are annotated as *"protein"*. Not included are proteins that are mentioned in the "Methods" section, e.g. restriction enzymes when creating an expression construct or those used as part of the protein purification process.

***Protein_state***

Properties of a protein are captured by the *"protein_state"*. This covers anything regarding its functional state, whether it carries a *"ptm"*, if it is bound to/void of/in complex with something, e.g. a component annotated as *"chemical"*. Also included are whether the protein is in its full-length native state or a variant/mutant or whether particular residues or a motif are conserved between related proteins.

***Protein_type***

Proteins are classed into different families/groups/classes based on their location in a cell or organism, or their function, or the process they are involved in. Terms capturing this are annotated as *"protein_type"* refers to a family/group/class of proteins. So a particular protein discussed in a publication is one example of the proteins covered by a whole family.

***PTM***

Post-translational modifications, *"ptm"*, cover any modification applied to a protein after creation of the main polypeptide chain. This covers modifications such as the formation of disulfide bridges, glycosylation, methylation, acetylation, ubiquitination, (auto)proteolytic cleavage and many more. These modifications rely on the creation/breakage of a covalent peptide bond. This excludes any covalent changes as part of a reaction mechanism, e.g. a covalent intermediate state which is covered by *"bond_interaction"*, and artificially introduced changes as part of an experiment.

***Residue_name***

This refers to a specific amino acid in the primary protein (a related protein for comparison) or a nucleic acid in an interaction partner such as DNA/RNA without giving a sequence position. This is distinct from an amino or nucleic acid serving as a *"chemical"* in a reaction mechanism of a protein.

***Residue_number***

If only the position in a sequence is given but the name of the residue, which can be either from a protein (primary or interaction partner or related), a peptide or a DNA/RNA, is not available then such a term is annotated as *"residue_number"*.

***Residue_name_number***

Residues in a sequence are uniquely identified by their name and position and therefore are annotated as *"residue_name_number"*. This applies to the primary protein of interest, interacting proteins or peptides and DNA/RNA (*"chemical"*). For proteins and peptides the residues in question are amino acids and for DNA/RNA they are nucleic acids.

***Residue_range***

A *"residue_range"* refers to a stretch of amino acid or nucleic acid residues, usually with a starting and ending position, or the number of residues spanning a particular stretch in a sequence. In cases where *"residue_name_number"* defines the start and end of the range, preference is given to *"residue_range"*.

***Site***

Any site of interest in the primary protein or in an interaction partner such as another protein or DNA/RNA is annotated *"site"*. This can be a binding site for a ligand, cofactor or metal, a protein-protein interaction point, specific residues involved in a mechanism.

***Species***

This entity type refers to specific species/strain the primary protein or a related protein originates from. However, this excludes species given in the "Methods" section.

***Structure_element***

Secondary structure elements are used to create more complex structural arrangements and division of a polypeptide into domains. Such structural features are annotated as *"structure_element"* in the primary protein as well as in any interaction partners such as other proteins, peptides or DNA/RNA. Also, for large complexes and molecular machines, a large number of very different polypeptides can be involved, each serving as a subunit of a larger assembly and therefore representing a *"structure_element"* to a larger *"complex_assembly"*.

***Taxonomy_domain***
This entity type captures organisms on a higher hierarchical level compared to *"species"*. Often a taxonomy term is used to refer to a specific organism ignoring the *"species"*. For example, "yeast" is generally assumed to refer to *"Saccharomyces cerevisiae"* and in our case the former was annotated as *"taxonomy_domain"* and the latter as *"species.* The exception is the word "human" which always refers to *"Homo sapiens"* and therefore is included in *"species.*

#### 1.2 Document sections

We restricted the manual annotation to the sections "Introduction", "Results", "Discussion", and "Conclusion" as well as tables and the captions for figures and tables. The named entities of interest were mainly found in the "Results" section and to a lesser extend in "Introduction", "Discussion", and "Conclusion", but almost no information could be gained from the "Methods" and the "References" and therefore these were excluded in the manual annotation. Also, any supplemental material was excluded as such contributions are usually in non-standardised formats which were not supported by the annotation tool. In the case of the iterative annotation with a semi-automated approach, annotations automatically added by the predictor to "Methods" and "References" were removed before using the data for the next round of training.

#### 1.3 Subspan annotation

In a number of publications we found authors used hand-crafted abbreviations. Such short-hands were split and annotated with separate entity types. Some examples are given below. "Hyp64IDA" represents a post-translational modification (*"ptm"*) for span fragment "Hyp64" and a "protein" for "IDA". Similarly, "H3K9me3" is also a post-translational modification *"ptm"* for the partial span "K9me3" whereas "H3" refers to a *"protein_type"*. "SceCD" refers to the entity type *"species"* as "Sce" stands for *"Saccharomyces cerevisiae"* and "CD" stands for "central domain" or "CD" covered by entity type *"structure_element"*. "Arg409HAESA" refers to the specific *"residue_name_number"* "Arg409" in *"protein"* "HAESA". And finally, some spans found for example *"residue_name_number"* like "Thr1OH" were only partially annotated, here "Thr1", as we were not interested in the specific group or atom in the residue that was involved in an interaction.

#### 1.4 Exclusion of non-specific spans

For the entity type *"chemical"* we also decided to limit annotations to specific chemical names and would ignore terms such as "moiety" or "group", if they referred to a part of a larger molecule. Also, more generally, any molecule that was not a protein was annotated as *"chemical"*. This could be nucleic acids, a peptide, some cofactor or ligand or some other small molecule or fragment interacting with the primary protein of interest.

##### 1.5 Usage of ontologies and controlled vocabularies

TeamTat allows linking entity types to ontologies and controlled vocabularies by defining a prefix, which can then be expanded with a reference to specific ID to ground a term. Although, we did not apply grounding in our project, we decided to define the prefix for a number of ontologies that are relevant for proteins and their structures: GO [1], [2], MESH [3], CHEBI [4], PR [5], SO [6], GENE [7] and DUMMY. DUMMY represents a placeholder for terms that were not found in any of the other ontologies or at the time of writing the authors were not aware of an ontology to hold these terms. Initially, annotators were also tasked in retrieving specific ontology IDs for each term. However, the time commitment for this task was not sustainable and therefore grounding for a specific ID was not applied after the initial set of annotations had been created. Instead, using ontology prefixes accelerated the annotation process. If a prefix for an ontology was set in TeamTat, an annotation could be automatically applied to all identical text spans in a publication when annotating manually.

#### 2 TeamTat User Guide

TeamTat `https://www.teamtat.org/about` is an open access, browser-based text annotation tool. In this user guide we describe how one can set up the tool for one's own project, how annotations can be added to a document, and how these annotations can be retrieved for downstream processes. To summarize the two different ways to operate TeamTat, as a project manager or annotator, we provide an overview for each of the participation options to emphasize the differences. Suppl Figure 1 displays the view an annotator will have when working on an annotation project and Suppl Figure 2 shows the more complex setup used by a project manager.

##### 2.1 Setting up TeamTat for an annotation project

###### 2.1.1 Starting a new project

A new project is started from TeamTat's landing page, which is given in Suppl Figure 3. If one has never used TeamTat before, then the first step is to click on *"Click here to Start"* at the very top, right-hand corner of the homepage. One is then asked to confirm to not be a robot in a pop-up window. After ticking the box and clicking on "Continue" one receives a unique "User" link, see Suppl Figure 4. This link represents one's unique access to the TeamTat tool and allows for the creation/management of annotation projects. As is stated in the image, the link can be shared with others, if multiple people are involved in managing a project. It is worth noting here, that there is a difference between a ”User” who manages a project and one who only annotates the text. The former has full access to all levels of the project, whereas the latter is restricted to only the publications they have been assigned to. Before continuing, one also needs to understand that if the project manager is also involved in annotating documents, then they should have a separate ”User” link for annotation work alongside the project manager link. Care should be taken to bookmark this unique user link to be able to easily return to one's projects.

To start a new project, one clicks on the button labelled ”New Project”, see Suppl Figure 5. Enter a descriptive project name. A project description can be given, but is not necessary. Generally, when using the unique user link, when clicking on ”Project” at the top of the page one is presented with all the projects one is involved in. An example of such a project list is given in Suppl Figure 6.

###### 2.1.2 Adding publications

TeamTat is fully integrated with PubMed and PubMedCentral, so publications can easily be added to a project by providing a list of IDs. For PubMed IDs (PMIDs) title and abstract will be retrieved for a given article, whereas for PubMedCentral IDs (PMCIDs), if a publication is open access, retrieval will cover the full-text including tables and figures and their captions. On the ”Documents” tab of the created project one clicks on the ”Add Documents” button, which will lead to a new window with upload and retrieval options Suppl Figure 7 and Suppl Figure 8. Please note that supplying an excessively long list of PMIDs or PMCIDs will fail and may even incapacitate the hosting server at NCBI. We recommend to keep the upload to ten publications at a time. Once all documents have been added to a project, one can navigate between them in a list via the ”Documents” tab Suppl Figure 9. The numerical part of a PMCID/PMID is used to create a unique document identifier in the table. Additionally, publication title and number of annotations for each document are shown in columns ”Title” and ”Annotation”, respectively. Columns ”Done”, ”Curatable” and ”Last Update” help monitoring when a publication is ready for downstream processes. There is an option to add documents by uploading from a local computer as text or PDF files. However, for our project, we did not use this option and cannot comment on how well this works.

###### 2.1.3 Adding team members

Annotators can be added to a project by re-using their name from a previous project or by adding them new. In order to keep things as anonymous as possible we decided to add all annotators by giving them non-identifiable names. Also, each annotator was given a personalized link to access the documents in the project and only the project manager knew which name and link corresponded to which annotator. Click on the button ”Add Anonymous Annotator”, add a non-identifiable name, bookmark the link and forward the link in an email to the annotator it is destined for. Setting a password is an option, but is not required for running a project. Once all team members have been given their personalized links, the project manager can assign documents. Please note again, if the project manager is also involved in the annotation process then they need to be given an annotator link as well. Also, please be aware that a project manager cannot have their manager link and annotator link open at the same time, even in separate browser windows, as annotations will always be created under the user ID that was last active. So if a project manager was using their manager link last and then wants to do annotation work they will have to switch links in order to have a consistent annotator for a document. Suppl Figure 10 shows a list of all members of an annotation project.

###### 2.1.4 Assigning publications to annotators

The tab "Assignments" allows the project manager to assign documents to individual annotators. This can be done manually by clicking on the circle in the document-annotator matrix or by using the "Random Assign" button, which will randomly assign publications to annotators, depending on a ratio specified by the project manager. Suppl Figure 11 shows the document-annotator matrix for a project aiming for double annotation of each document.

###### 2.1.5 Defining entity types

The project manager is also required to define and add entity types to a project. Using the tab "Types" this can be set up. Here, the project manager first clicks on the button "New Entity Type" to be prompted with a form which asks for "Name" and "Prefix" for the new entity type. As was explained in the main text of our publication, the "Prefix" enables linking of an entity type to an ontology or controlled vocabulary. Colors can be chosen for the different entity types. The colors can be personalized by each team member, including the annotators, and only applies to each individual link. In case an annotator may be colorblind this will allow them to choose a set of colors they are able to differentiate. Suppl Figure 12 gives the input form to create a new entity type and Suppl Figure 13 shows an example list of defined entity types.

###### 2.1.6 Opening/Closing an annotation round

An annotation round is opened/closed for all annotators and their assigned documents by clicking on the button "Start Round" (see Suppl Figure 14). The project manager has to choose between two options. Using the "Individual Round", as was done for our project, means that the different annotators work independently and cannot see the annotation results of each other while annotating. Once an annotation round has been closed annotations are exchanged so annotators can see each other's work. For the "Collaborative Round" annotators can see each other's annotations and can therefore work together when resolving annotation disagreements. While an annotation round is open, the project manager can see how many annotations for each entity type have been found by the annotators and how many unique text spans they represent. Once all documents have had their status set by the annotators an annotation round can be closed. Closing a round will automatically set TeamTat to calculate inter-annotator agreement for each document. Suppl Figure 15 gives a shortened list of annotations found for different entity types. An example of calculated inter-annotator agreement can be found in Suppl Figure 16.

#### 2.2 Using TeamTat when annotating

As mentioned above, the annotator view in TeamTat has limited options available which are only focused on the annotation process and not the project management. Once a project manager has assigned all the documents to the annotators and has opened an annotation round, an annotator can access the documents through their personalized link. Suppl Figure 17 displays an example of a document list an annotator finds in their project. By clicking on the document identifier one can select and open a document to work on (see Suppl Figure 18).

##### 2.2.1 Creating an annotation

There are multiple ways how an annotator can create an annotation. Below are examples for how one may want to annotate a text span with an entity type. Importantly, TeamTat does not have an "undo" function. If annotations are created/removed in error then they have to be manually removed/added to remedy the mistake. Also, it is good practice to reload the browser page to ensure that the autosave function of the tool has indeed carried all changes over to the document.

A combination of versions 2-4 has proven to be the most time efficient and user friendly way to annotate a text. Version 5 is a very good way when resolving annotation conflicts and cleaning and consolidating annotations.

Version 1 (Suppl Figure 19)

1. select entity type from drop-down menu
2. mark text span

Version 2 (Suppl Figure 20)

1. mark text span
2. click on text span
3. edit text span in pop-up window/delete annotation
4. click on "Update" button

Version 3 (Suppl Figure 21)

1. click on magnifying glass next to an item in the list/tab "Annotations"

2. text will move to focus on selected entity
3. the entity of interest is high-lighted by a red underline
4. edit text span in the pop-up window/delete annotation
5. click on "Update" button

Version 4 (Suppl Figure 22)
Annotating the same text span throughout an entire publication requires multiple clicks for each instance. However, by linking an annotation to an ontology using the "Prefix" one can annotate all instances of the same text span by following the instructions below. Note, never use "Update all mentions with the same concept ID" unless there is a fine-grained selection/hierarchy of ontologies in place. If "Concept ID" is not set, then using the "Update all mentions with the same concept ID" option will corrupt annotations.

1. either click on marked text span or select from list and click on magnifying glass
2. edit text span in the pop-up window
3. click on "Annotate each instance of this mention text"
4. if needed also set "Case sensitive match"
5. "Match whole word only" is set by default

Version 5 (Suppl Figure 23 and Suppl Figure 24)

1. highlight text span, sentence or paragraph containing annotations
2. edit existing annotations from list in pop-up window
3. do not click on "+Create New Annotation"

Note, this will create a new annotation for the entire highlighted text, which one does not want to create, but at the same time also brings any existing annotation into a new pop-up window as a list. One can then work their way through all entries in the list, if necessary. In particular, for removing annotations this is a very convenient way to reduce the number of clicks.

###### 2.2.2 Flagging documents ready for curation

Each document has two sliders at the top that allow an annotator to set the status of the document as "Curatable" and "Done" which will be reflected in the document list for annotator and project manager (see Suppl Figure 25). These flags tell the project manager when the different documents are ready and they can close an annotation round.

#### 2.3 Curating annotations

After closing an annotation round in our project, the curation and consolidation of annotations was left to the project manager with the lead biocurator serving as a proof-reader. Inter-annotator agreement was calculated at this stage, as was already mentioned above, and all the annotations of the different annotators were combined into their respective documents. Suppl Figure 26 shows how the text will be displayed with all the annotations combined. The project manager was then required to go through all the annotations to either accept them, as they were in full agreement between the two annotators, apply the necessary fixes to produce an agreement or add annotations, if they were missing. In order to keep track of changes and be able to calculate inter-annotator agreement during the different curation rounds, the project manager used the personalized annotator links. Suppl Figure 27 gives an idea of how the different versions to annotate text described above can be used to work through the list of annotations. The side panel that holds a list of all the annotations in the text also indicates which require attention by the curator. We also found that the "Skip" button stayed active once it had been clicked and resulted in annotations worked on afterwards were also skipped.

#### 2.4 Downloading annotated documents

Once the annotation project has reached desired maturity, the documents can be downloaded in a variety of ways. Individual documents can be downloaded by selecting a document from the the list in the "Documents" tab, clicking on it and then using the "Download" button at the top choosing between BioC XML or JSON as format. Using the "Version" button at the top of the document one is able to download a specific version of the document, depending on the annotation rounds carried out (see Suppl Figure 28). Only the project manager is able to download all documents in a project at once. This can be accomplished by using the "Download" button on the project overview page (see Suppl Figure 29). After selecting the version of the project one wants to download, a compressed folder of all the documents with their annotations in BioC XML is created. The "BioCXML" version contains the publication cut into paragraphs and set within XML tags <passage> Each paragraph has its corresponding annotations included within the tags. Individual annotations are surrounded by the tag <annotation id="xxxx> which also includes a unique, non-changeable ID for the annotation. The text span covered by the annotation as well as the entire text of the paragraph is enclosed within the tags <text>. Offsets based on character counts allow to determine the start and end of an annotation and where it is located in the document. The "PubAnnotatorJSON" version contains a JSON dictionary with keys "sourceid", "sourcedb", "project", "target", "text" and "denotations". The plain text as single string is found under "text". The annotations are collected as a list of dictionaries under "denotations". The individual dictionaries contain the following keys "span", "obj", "id". The key "span" itself contains a dictionary with keys "begin" and "end" giving the the start and end position of the text span with respect to the character offset for the text not considering individual paragraphs. The key "obj" contains a string crafted from the entity type name, the set "Prefix" to link to an ontology, the annotator and a time stamp. The key "id" refers to a unique, non-changable ID for the annotation. The annotated text span itself is not included. The document and text and the found annotations are not directly linked.

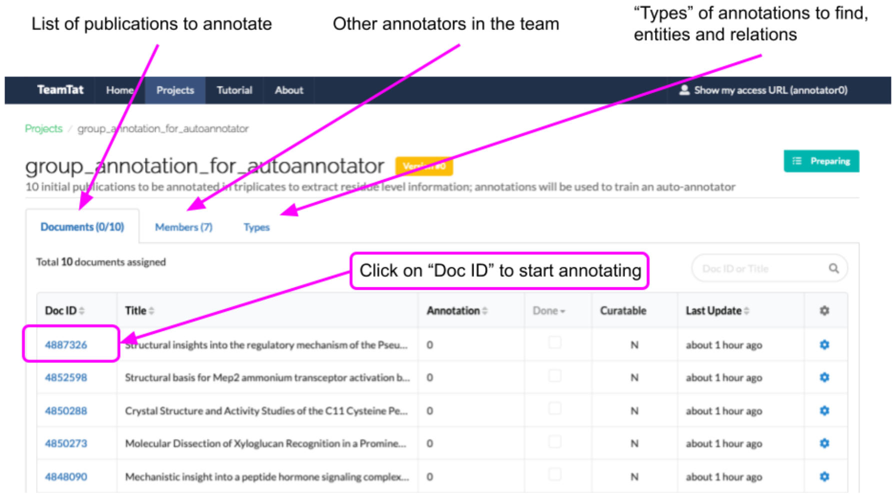


Figure 1: Annotator view for TeamTat

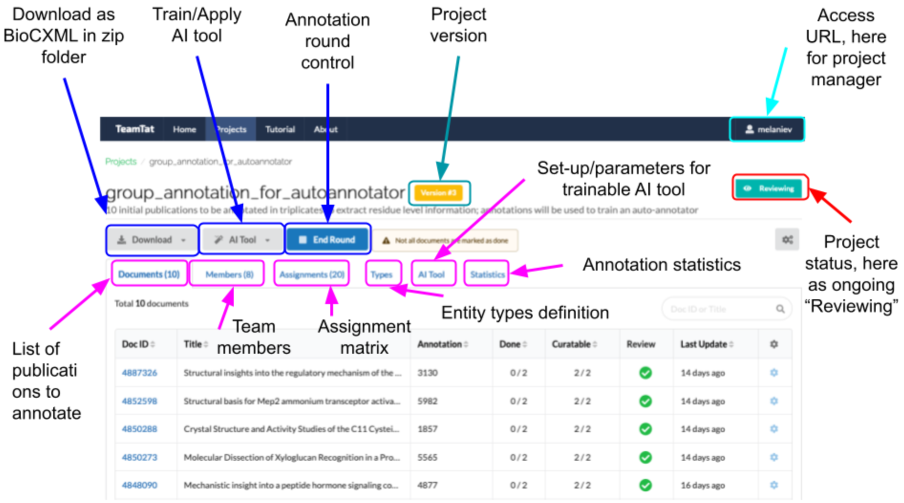


Figure 2: Project manager view for TeamTat

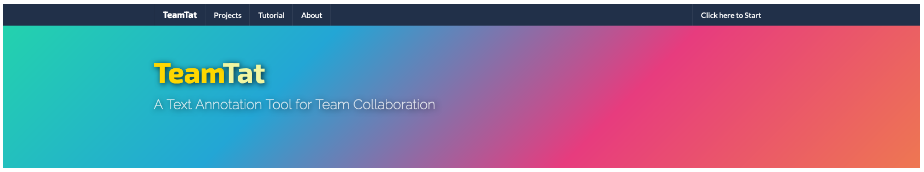


Figure 3: Arriving on TeamTat's homepage

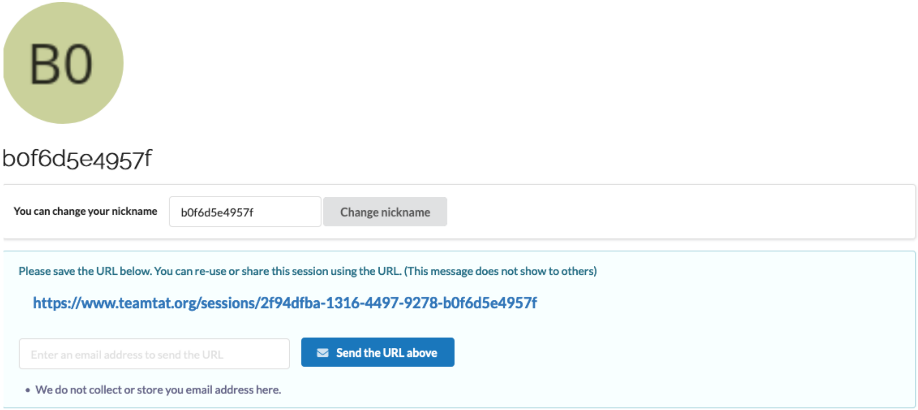


Figure 4: Receiving unique user link

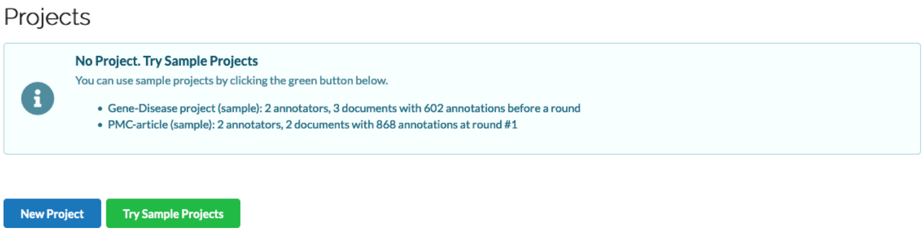


Figure 5: Creating a new project

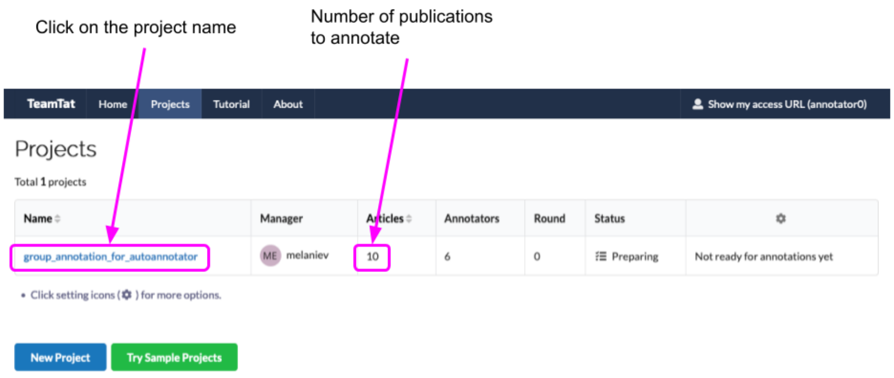


Figure 6: List of projects one is part of

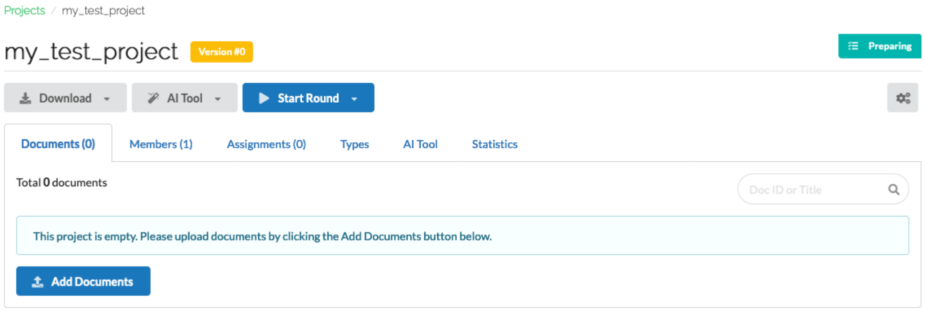


Figure 7: Input form to add documents to a project

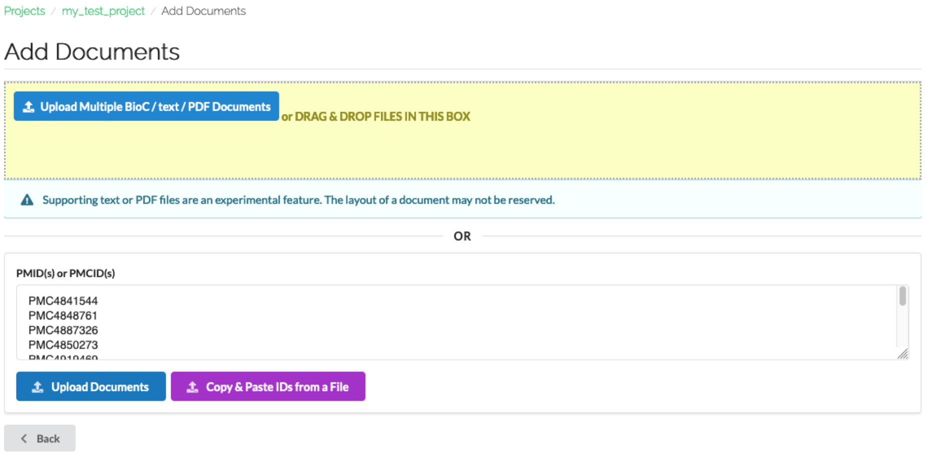


Figure 8: Adding publications by providing a list of PMCIDs

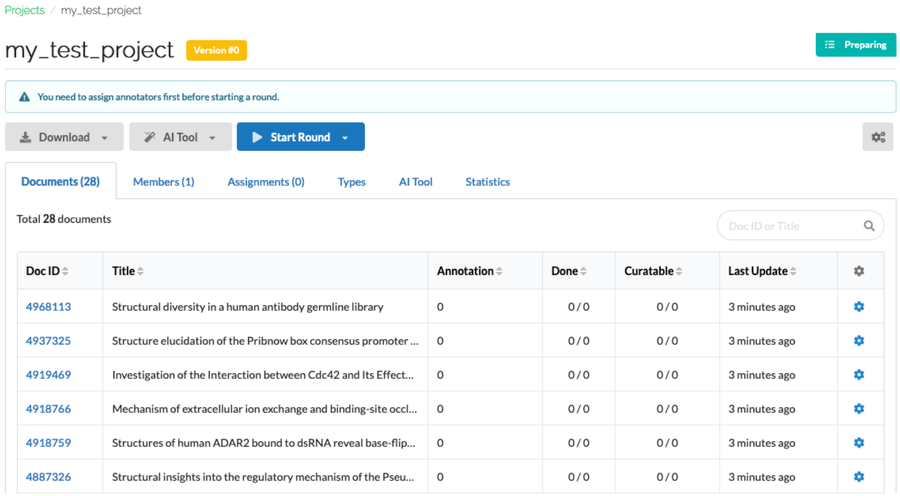


Figure 9: Example of a document list for a project

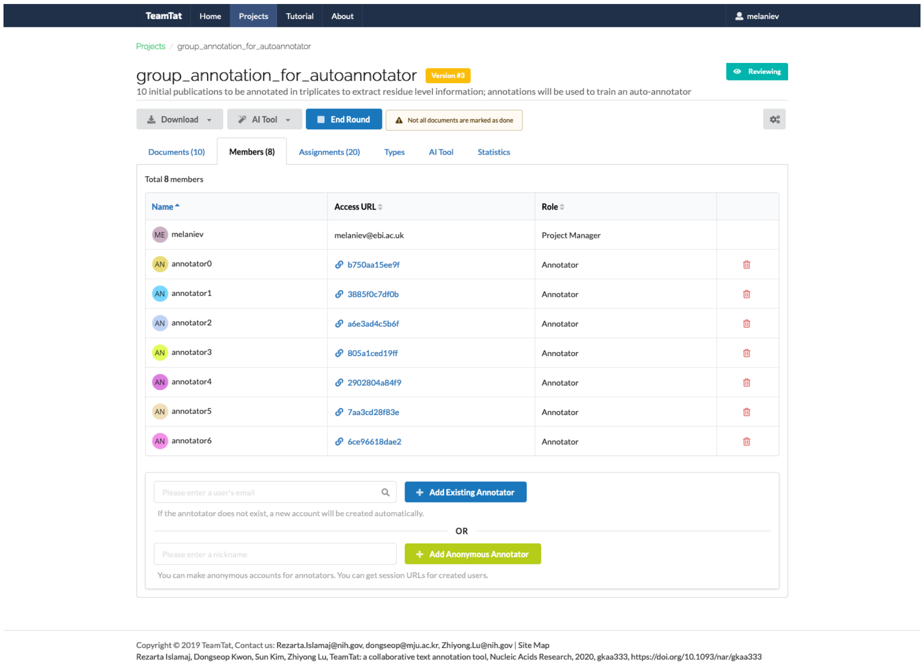


Figure 10: List of team members in an annotation project

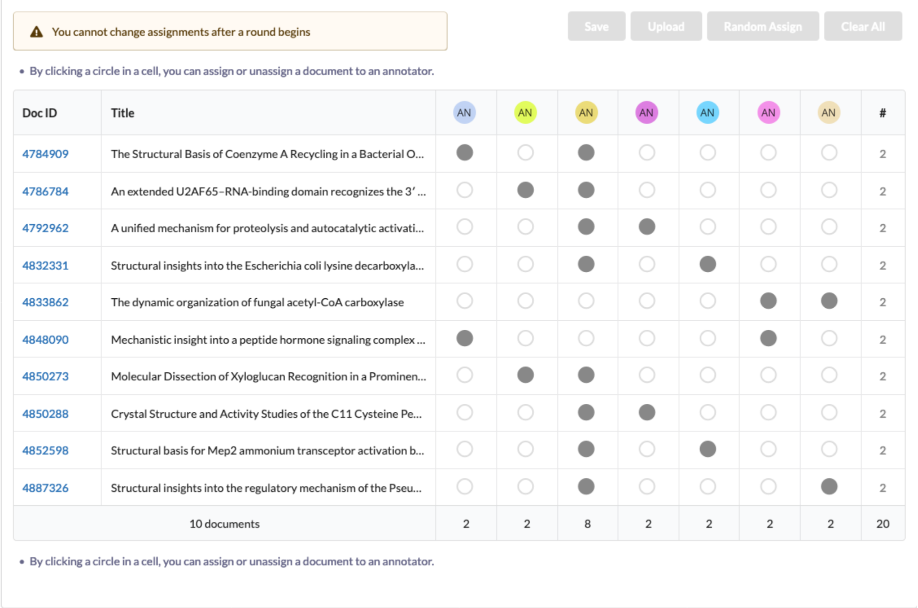


Figure 11: Document-annotator matrix for a project aiming for double annotation of each document

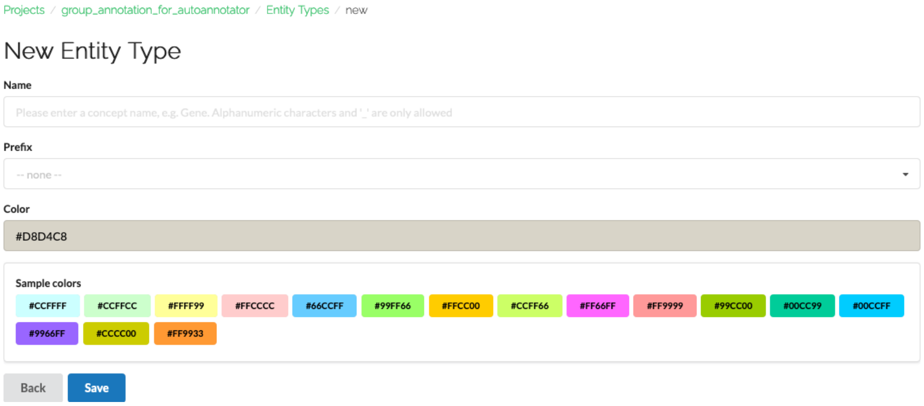


Figure 12: Input form to create a new entity type

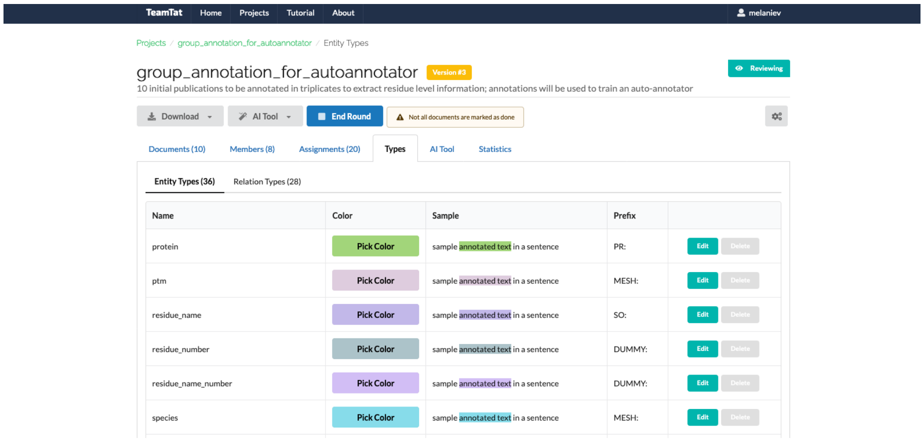


Figure 13: Example list of different entity types

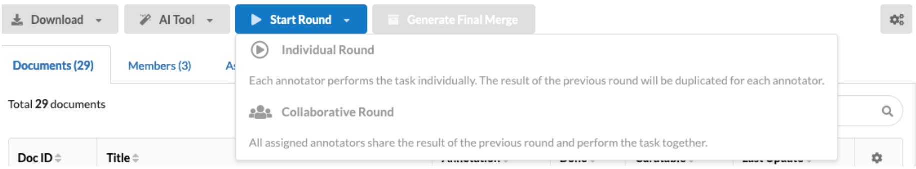


Figure 14: Opening/closing of an annotation round

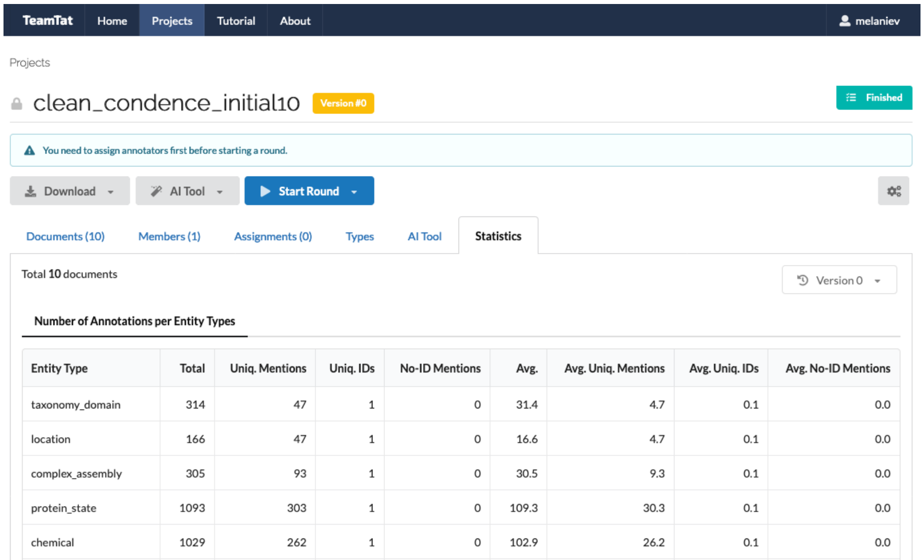


| Entity Type | Total | Uniq. Mentions | Uniq. IDs | No-ID Mentions | Avg. | Avg. Uniq. Mentions | Avg. Uniq. IDs | Avg. No-ID Mentions |
|---|---|---|---|---|---|---|---|---|
| taxonomy_domain | 314 | 47 | 1 | 0 | 31.4 | 4.7 | 0.1 | 0.0 |
| location | 166 | 47 | 1 | 0 | 16.6 | 4.7 | 0.1 | 0.0 |
| complex_assembly | 305 | 93 | 1 | 0 | 30.5 | 9.3 | 0.1 | 0.0 |
| protein_state | 1093 | 303 | 1 | 0 | 109.3 | 30.3 | 0.1 | 0.0 |
| chemical | 1029 | 262 | 1 | 0 | 102.9 | 26.2 | 0.1 | 0.0 |

Figure 15: Example of annotations for different entity types and their raw counts

| Doc ID | # | FA | CA | TA | PA | DA | SN | FA (%) | CA (%) | TA (%) | PA (%) | DA (%) | SN (%) |
|---|---|---|---|---|---|---|---|---|---|---|---|---|---|
| 4784909 | 1126 | 1102 | 4 | 0 | 1 | 7 | 12 | 97.87 | 0.36 | 0.0 | 0.09 | 0.62 | 1.07 |
| 4786784 | 1961 | 1909 | 9 | 0 | 7 | 12 | 24 | 97.35 | 0.46 | 0.0 | 0.36 | 0.61 | 1.22 |
| 4792962 | 1806 | 1771 | 0 | 0 | 7 | 7 | 21 | 98.06 | 0.0 | 0.0 | 0.39 | 0.39 | 1.16 |
| 4832331 | 997 | 985 | 2 | 0 | 1 | 0 | 9 | 98.8 | 0.2 | 0.0 | 0.1 | 0.0 | 0.9 |
| 4833862 | 1432 | 1387 | 3 | 0 | 6 | 18 | 18 | 96.86 | 0.21 | 0.0 | 0.42 | 1.26 | 1.26 |
| 4848090 | 1360 | 1329 | 0 | 0 | 2 | 8 | 21 | 97.72 | 0.0 | 0.0 | 0.15 | 0.59 | 1.54 |
| 4850273 | 1492 | 1335 | 10 | 0 | 2 | 65 | 80 | 89.48 | 0.67 | 0.0 | 0.13 | 4.36 | 5.36 |
| 4850288 | 928 | 916 | 0 | 0 | 2 | 4 | 6 | 98.71 | 0.0 | 0.0 | 0.22 | 0.43 | 0.65 |
| 4852598 | 1679 | 1405 | 10 | 4 | 18 | 189 | 53 | 83.68 | 0.6 | 0.24 | 1.07 | 11.26 | 3.16 |
| 4887326 | 1297 | 1273 | 2 | 1 | 2 | 15 | 4 | 98.15 | 0.15 | 0.08 | 0.15 | 1.16 | 0.31 |
| Total | 14078 | 13412 | 40 | 5 | 48 | 325 | 248 | 95.27 | 0.28 | 0.04 | 0.34 | 2.31 | 1.76 |

- **FA - Full Agree:** same type, concept ID and text span
- **CA - Concept Agree:** same concept ID and text span, but different types
- **TA - Type Agree:** same type and text span, but different concept IDs
- **PA - Partial Agree:** same type and concept ID for overlapping text
- **DA - Disagree:** different types, concept IDs or text spans
- **SN - Single:** text annotated by only some of annotators

Figure 16: Example of calculated inter-annotator agreement

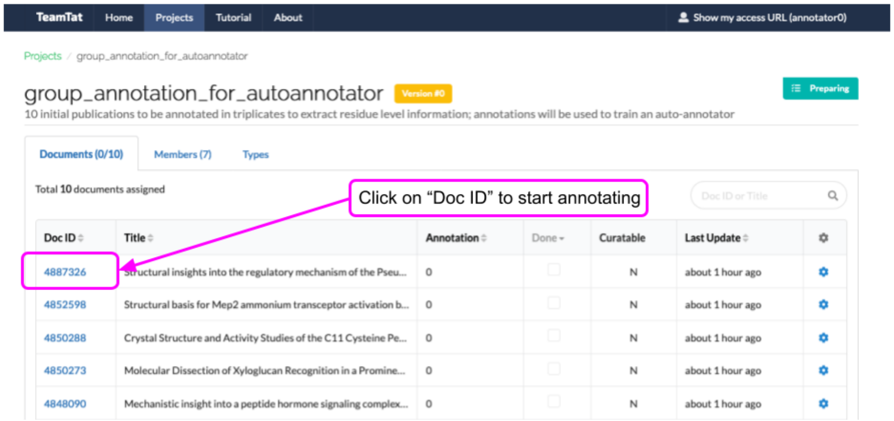


Figure 17: Example of a document list for an annotator

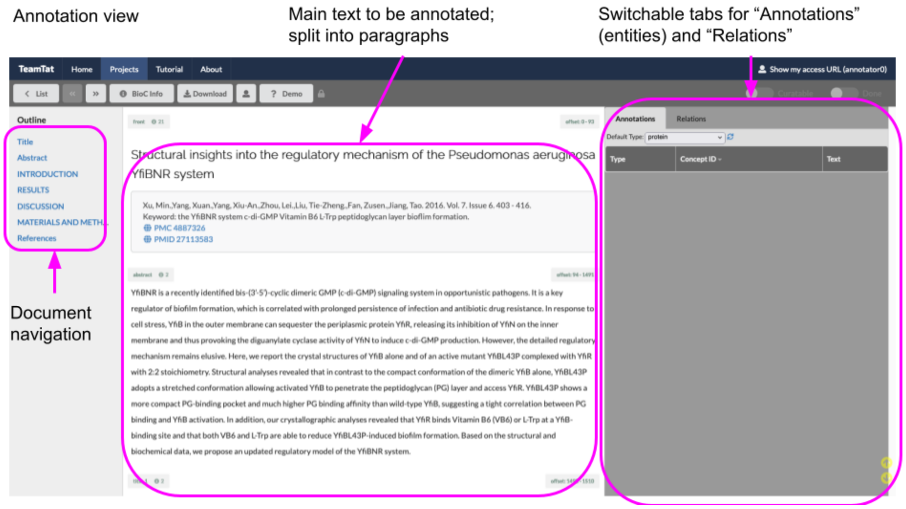


Figure 18: View when opening a document for annotation

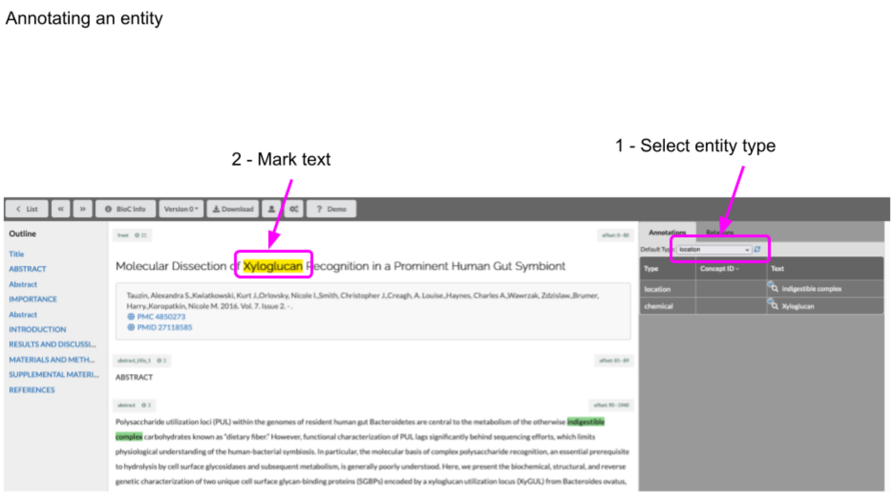


Figure 19: Creating a new annotation, version 1

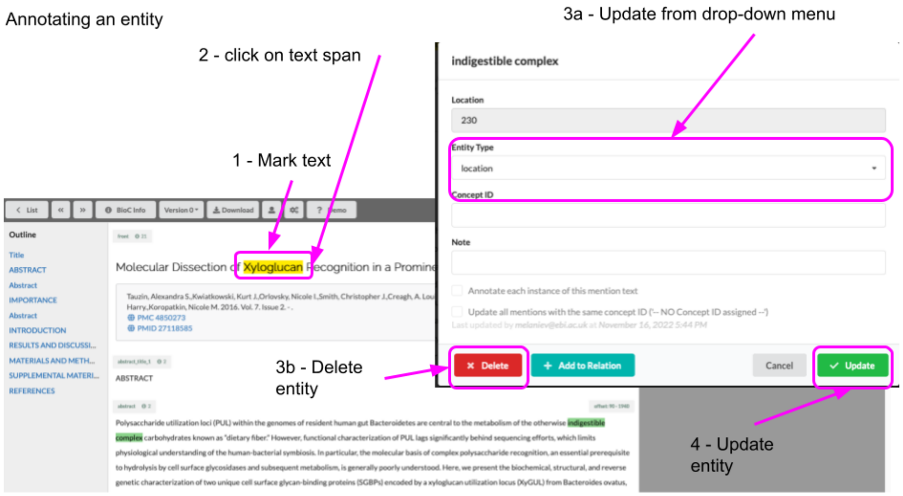


Figure 20: Creating a new annotation, version 2

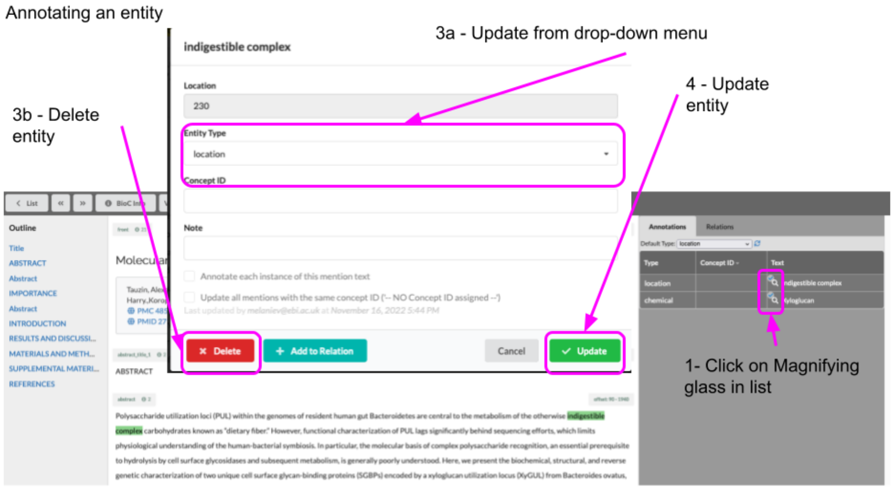


Figure 21: Creating a new annotation, version 3

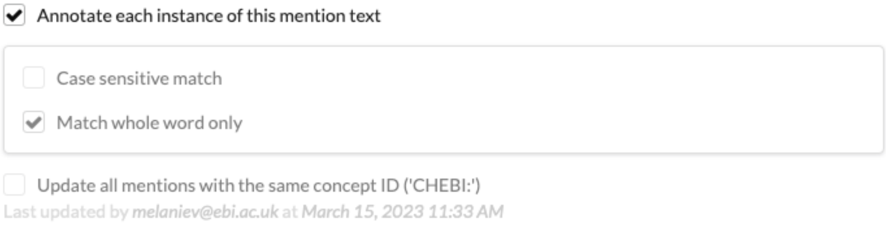


Figure 22: Creating a new annotation, version 4

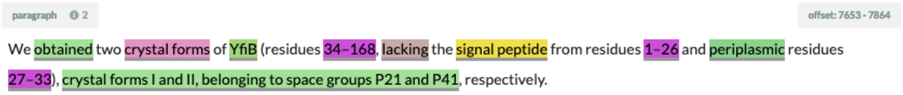


Figure 23: Selecting text with multiple annotations

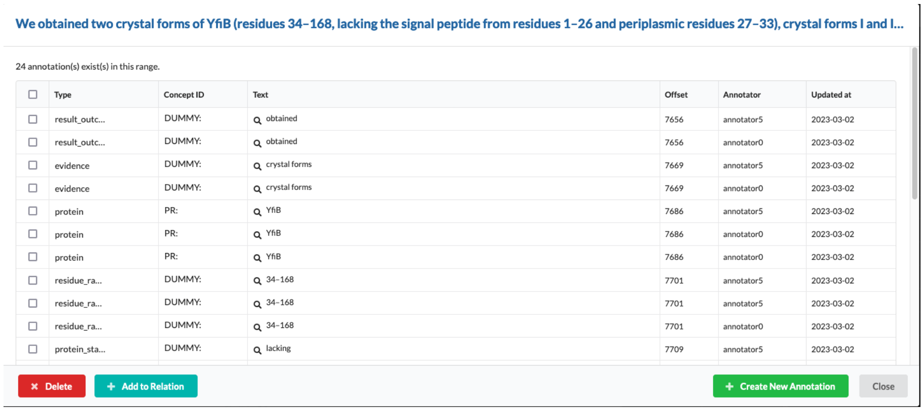


Figure 24: Updating multiple annotations at the same time

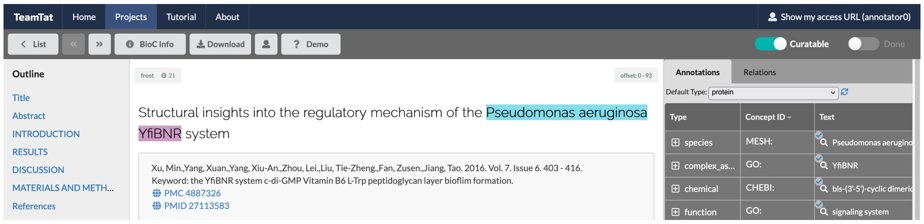


Figure 25: Example showing sliders to set document status

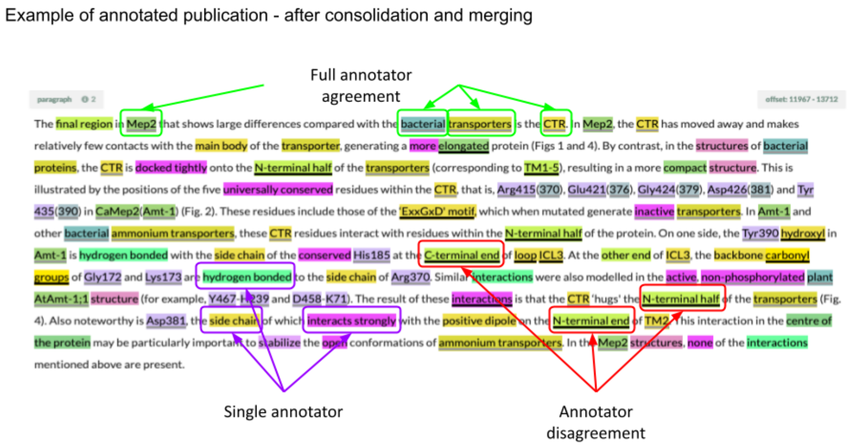


Figure 26: Example text showing the different outcomes when annotations of two annotators are combined after closing an annotation round

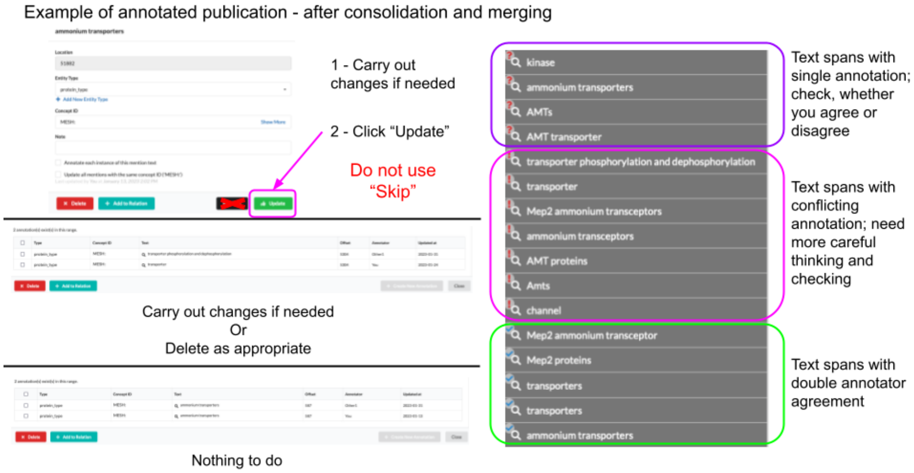


Figure 27: Different ways to curate annotations

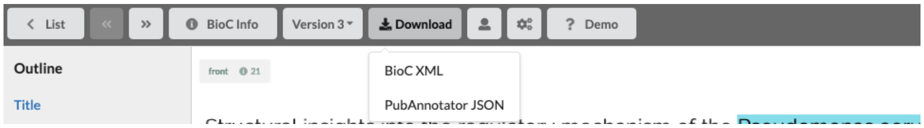


Figure 28: Selecting download of a single document

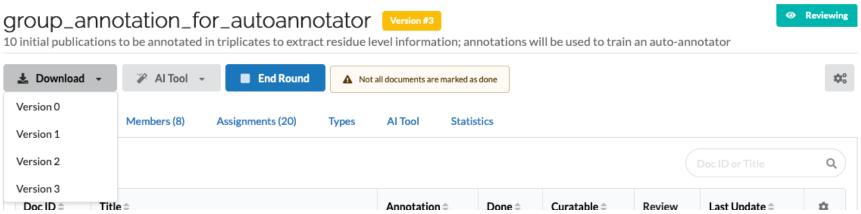


Figure 29: Downloading all documents of a project

| Entity type | Example |
|---|---|
| bond_interaction | H-bond, hydrogen bond network, hydrophobic interaction, salt bridge |
| chemical | NADH, ATP, Zn2+, RNA, acetyl CoA |
| complex_assembly | Tdp2-DNA, BRCA2:RAD51, TTD–PHD–H3K9me3, UHRF1–DNMT1 |
| evidence | KD, structure, root-mean-square deviation, chromatogram, electron density |
| experimental_method | size exclusion chromatography, mass spectrometry analysis, sequence alignment, analytical ultracentrifugation, single-residue mutation |
| gene | ectC, Synpcc7942_2462, At2g21370, YOR006c, nep1, nadA, nadR |
| mutant | YfiBL43P, H7A, MC58-Δ1843, ΔNadR, mep1-3Δ, 449-485Δ |
| oligomeric_state | monomer, dimer, trimer, monomeric, dimeric, heterodimer |
| protein | YfiB, YfiR, YfiN, HAESA, SERK1, SNF1, BRCA1 |
| protein_state | phosphorylated, non-catalytic, highly conserved, full-length, biodegradative, apo, ppGpp-free |
| protein_type | cyclase, lipase, hydrolase, kinase, phosphatase |
| ptm | glycosylation, phosphorylation, methylation, acetylation, disulfide bridges, propeptide cleavage, K9me3, Diph699, pThr160 |
| residue_name | alanine, proline, tyrosine, Glu, Lys, His, adenosine, A, guanosine, G |
| residue_number | 123 |
| residue_name_number | Ala123, A123, X1, X2, U11, mAsp272, hAsp262, glutamate at 474, G-1 |
| residue_range | 34-52, D21-K26, Arg120 until Ser122, Gln5 to Ser122, 15 residues, three amino acids |
| site | hydrophobic cleft, HAP binding site, Trp binding pocket, haem-binding region, FXF-motif-binding site, active site |
| species | *E. coli, Pseudomonas aeruginosa, Sphingopyxis alaskensis, S. alaskensis, Sa* |
| structure_element | beta-sheets, cupin barrel, $\alpha$-helices, metal-binding motif, antiparallel $\beta$-strands |
| taxonomy_domain | Bacteria, nitrifying archaeon, eukaryotic, *Xenopus*, murin, mammalian |

Table 1: Entity types with examples.
